# Unfolded protein-independent IRE1 activation contributes to multifaceted developmental processes in Arabidopsis

**DOI:** 10.1101/651182

**Authors:** Kei-ichiro Mishiba, Yuji Iwata, Tomofumi Mochizuki, Atsushi Matsumura, Nanami Nishioka, Rikako Hirata, Nozomu Koizumi

## Abstract

As an initial step for the unfolded protein response (UPR) pathway, the luminal domain of inositol requiring enzyme 1 (IRE1) senses unfolded proteins in the endoplasmic reticulum (ER). Recent findings in yeast and metazoans suggest alternative IRE1 activation without the sensor domain, although its mechanism and physiological significance remain to be elucidated. In Arabidopsis, the *IRE1A* and *IRE1B* double mutant (*ire1a/b*) is unable to activate cytoplasmic splicing of *bZIP60* mRNA and regulated IRE1-dependent decay (RIDD) under ER stress, while the mutant does not exhibit severe developmental defects and is fertile under non-stress conditions. In this study, we focused on a third Arabidopsis *IRE1* gene, designated as *IRE1C*, whose product lacks a sensor domain. We found that even though *ire1c* and *ire1a/c* mutants did not exhibit defective *bZIP60* splicing and RIDD under ER stress, the *ire1a/b/c* triple mutant is lethal. Heterozygous *IRE1C* (*ire1c*/+) mutation in the *ire1a/b* mutants resulted in growth defects and reduction of the number of pollen grains. Genetic analysis revealed that *IRE1C* is required for male gametophyte development in the *ire1a/b* mutant background. Expression of a mutant form of IRE1B that lacks the luminal sensor domain (ΔLD) in the *ire1a/b* mutant did not complement defects in ER stress-dependent *bZIP60* splicing and RIDD. Nevertheless, expression of ΔLD complemented a developmental defect in the male gametophyte in *ire1a/b/c* haplotype. *In vivo*, the ΔLD protein was activated by glycerol treatment that increases the composition of saturated lipid and was able to activate RIDD but not *bZIP60* splicing. Phenotypes of *IRE1B* mutants lacking the sensor domain produced by CRISPR/Cas9-mediated gene editing in the *ire1a/c* mutant background were essentially same as those of ΔLD-expressing *ire1a/b* mutant. These observations suggest that IRE1 contributes to plant development, especially male gametogenesis, using an alternative activation mechanism that bypasses the unfolded protein-sensing luminal domain.

## Introduction

The endoplasmic reticulum (ER) in eukaryotes copes with an accumulation of unfolded proteins by activating the unfolded protein response (UPR), which increases protein folding capacity and attenuates protein synthesis in the ER (Walter & Ron, 2011). Inositol-requiring enzyme 1 (IRE1) is the primary transducer of the UPR. IRE1 consists of an N-terminal sensor domain facing the ER lumen, a single transmembrane helix (TMH) embedded in the ER membrane, and kinase and ribonuclease (RNase) domains at its C-terminus on the cytosolic side (Nikawa & Yamashita, 1992; Sidrauski & Walter, 1997). Under ER stress, IRE1 senses unfolded proteins in the ER, which causes autophosphorylation of IRE1 to exert RNase activity for cytoplasmic splicing. Targets of the cytoplasmic splicing are mRNAs encoding UPR-specific transcription factors, such as HAC1 in yeasts (Sidrauski & Walter, 1997), XBP1 in metazoans (Yoshida *et al*, 2001) and bZIP60 in Arabidopsis (Deng *et al*, 2011; Nagashima *et al*, 2011). Activated IRE1 also degrades mRNAs encoding secretory pathway proteins, designated as the Regulated IRE1-Dependent Decay (RIDD) of mRNAs in fission yeast (Kimmig et al, 2012), metazoans (Hollien & Weissman, 2006; Iqbal *et al*, 2008; Han *et al*, 2009; Hollien *et al*, 2009) and plants (Mishiba *et al*, 2013; Hayashi *et al*, 2016). Although distinct catalytic mechanisms between cytoplasmic splicing and RIDD has been reported (Tam *et al*, 2014), how IRE1 outputs these two modules during physiological and developmental processes is still unclear (Maurel *et al*, 2014).

Even though IRE1-deficient mice (Zhang *et al*, 2005) and flies (Ryoo *et al*, 2013) cause embryonic lethality, IRE1-deficient yeast (Nikawa & Yamashita, 1992; Kimmig *et al*, 2012) and worms (Shen *et al*, 2001) are viable. In plants, Arabidopsis IRE1A- and IRE1B-defective mutants do not exhibit severe developmental phenotypes under normal conditions (Nagashima *et al*, 2011; Chen & Brandizzi, 2011), whereas rice homozygotes that express kinase-defective IRE1 is lethal (Wakasa *et al*, 2012; note that rice has one *IRE1* gene). This different consequence of IRE1 mutation in phenotypes between Arabidopsis and rice prompted us to investigate the degree of contribution that IRE1 makes to plant development.

In recent years, activation of IRE1 caused by lipid perturbation was observed in yeast (Promlek *et al*, 2011) and mouse cells (Volmer *et al*, 2013). This IRE1 activation does not require sensing of unfolded proteins by the luminal domain of IRE1, but does require an amphipathic helix (AH) adjacent to the transmembrane helix (TMH) to sense ER membrane aberrancies (Halbleib *et al*, 2017). Although physiological functions of lipid-dependent IRE1 activation is less well known, it has been presumed that the unfolded protein-independent mechanism allow cells to prospectively adopt their ER folding capacity (Volmer & Ron, 2015). For instance, mutant worms with decreased membrane phospholipid desaturation activate IRE1 without promoting unfolded protein aggregates (Hou *et al*, 2014). However, there are no studies directly addressing the importance of the unfolded protein-independent IRE1 activation in developmental processes in multicellular organisms.

In this report, we investigated the contribution of IRE1 lacking its sensor domain to Arabidopsis development. We found that a third Arabidopsis *IRE1* gene, encoding sensor domain-lacking IRE1, is functional and that the triple mutant of the three *IRE1* (*IRE1A-C*) genes is lethal. Our analyses with plants that express mutant IRE1B proteins without the sensor domain suggest contribution of unfolded protein-independent IRE1 activation to multifaceted developmental processes in Arabidopsis.

## Results

### Loss of function of IRE1C does not alter ER stress response

In addition to the *IRE1A* and *IRE1B* genes, Arabidopsis contains an *IRE1-like* gene (AT3G11870; designated as *IRE1C* hereafter), whose product lacks a sensor domain (Fig. 1A). The sensor domain-lacking IRE1 was also found in some other Brassicaceae species, such as *Camelina sativa*, and phylogenetic analysis showed that the IRE1C forms an independent cluster from IRE1A and IRE1B groups in dicotyledonous plants (Fig. 1B). A T-DNA insertion mutant of *ire1c* (SALK_204405; Fig. S1A) and *ire1a ire1c* (designated as *ire1a/c* hereafter) double mutants did not exhibit any visible phenotypic alterations in normal growth conditions (Fig. 1C). Consistent with the previous studies (Nagashima *et al*, 2011; Mishiba *et al*, 2013), susceptibility to ER stress inducer, dithiothreitol (DTT) was more apparent in *ire1a/b* mutant than those in WT, *ire1a*, and *ire1b* mutants (Fig. 1D). The susceptibility to DTT in *ire1c* and *ire1a/c* mutants was same as that in WT (Fig. 1D). To detect *bZIP60* splicing and RIDD in the *IRE1* mutants under ER stress, expressions of *BiP3* and *PR-4* mRNA, which are the typical targets of bZIP60 (Iwata & Koizumi, 2005) and RIDD (Mishiba *et al*, 2013), respectively, were analyzed by northern blotting. Up-regulation of *BiP3* mRNA and down-regulation of *PR-4* mRNA by tunicamycin (Tm) treatment was observed in *ire1c* and *ire1a/c* mutants as well as WT, *ire1a*, and *ire1b* mutants, but not in *ire1a/b* mutant (Fig. 1E). These results indicate that IRE1C, which lacks a sensor domain, does not contribute to ER stress response in Arabidopsis.

**Figure 1.**
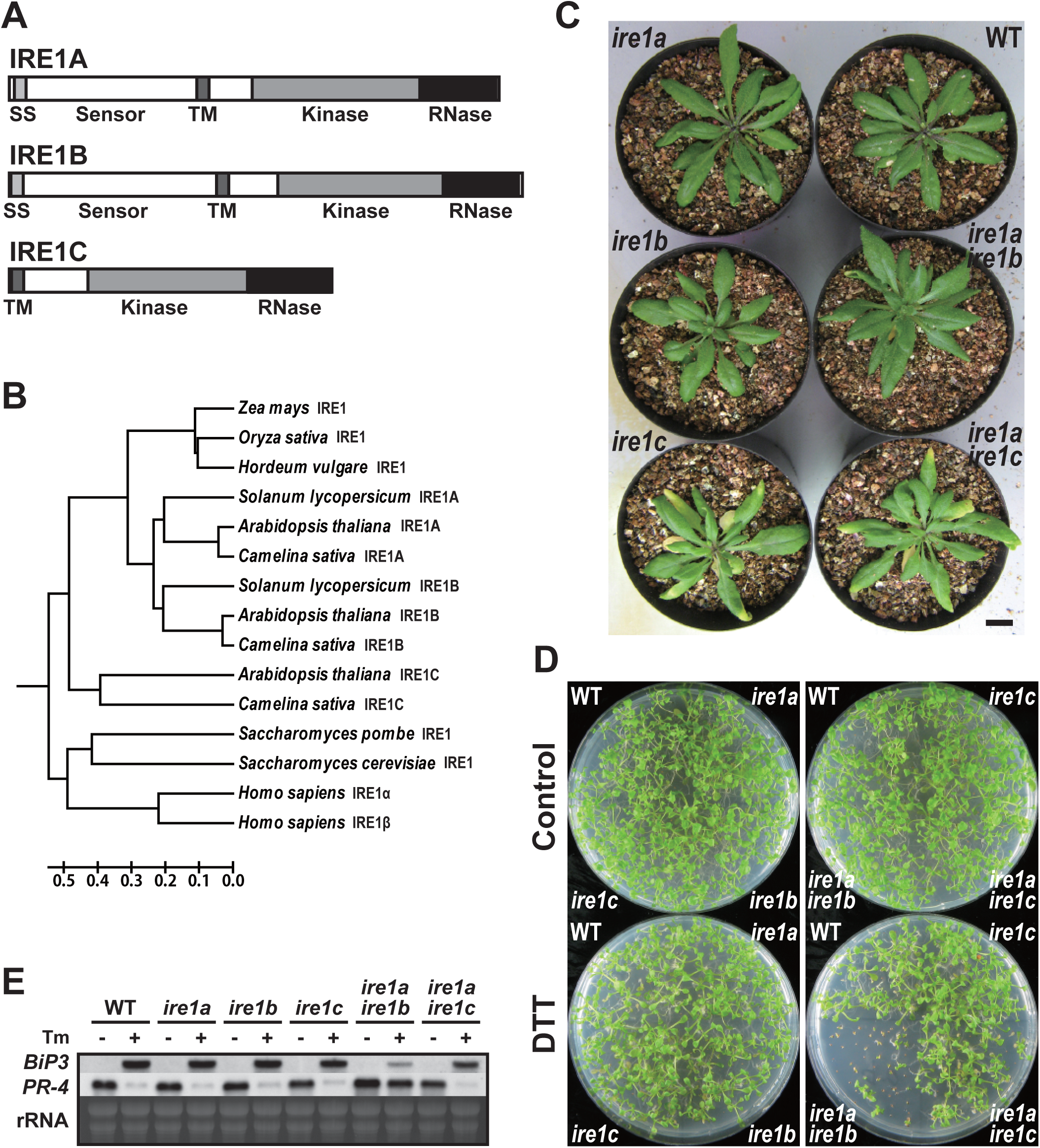
Arabidopsis IRE1C does not contribute to the unfolded protein response. (A) Structures of Arabidopsis IRE1A, IRE1B and IRE1C proteins. SS, signal sequence; TM, transmembrane domain. (B) Phylogenetic tree of IRE1-like proteins from mammals (*Homo sapiens*), fungi (*Saccharomyces cerevisiae* and *Saccharomyces pombe*) and plants was constructed by UPGMA using MEGA 6 (Tamura *et al*, 2013). The tree is drawn to scale, with branch lengths in the same units as those of the evolutionary distances used to infer the phylogenetic tree. The evolutionary distances were computed using the Poisson correction method and are in the units of the number of amino acid substitutions per site. (C) Wild-type (WT) and *ire1* mutant plants 40 days after germination (DAG). Bar = 10 mm. (D) DTT sensitivity of the *ire1* mutants. Seedlings at 15 DAG of the indicated lines were treated with or without 1 mM DTT. (E) RNA blot analysis of *BiP3* and *PR-4* in WT and *ire1* mutants. Seedlings at 10 DAG were treated with (+) or without (−) 5 mg/l tunicamycin (Tm) for 5 h.

### A triple mutant of *IRE1A*, *IRE1B*, and *IRE1C* is lethal

We tried to produce *ire1a/b/c* triple mutant by crossing *ire1a/b* with *ire1c* mutants. We obtained three F_2_ plants heterozygous for *ire1c* (designated as *ire1c*/+) and homozygous for *ire1a* and *ire1b*. Genotyping of their self-pollinated progenies showed that no plants homozygous for *ire1c* were obtained among 108 plants analyzed (Table 1, Fig. S1B). Additionally, unexpected segregation of homozygotes and heterozygotes for *IRE1C* (+/+: *ire1c*/+ = 1.0:0.3) was observed. These results indicate that *ire1a/b/c* triple mutant is lethal. The *ire1a/b ire1c*/+ plants exhibited growth retardation (Fig. 2A) and reduced seed set (Figs. 2B, 2C) compared to the *ire1a/b* +/+ siblings. Pollen development was especially impaired in the *ire1a/b ire1c*/+ plants, whereas *ire1c* and *ire1a/c* mutants did not affect pollen development and seed set (Fig. 2C). No transmission of the *ire1c* allele through male gametophyte were shown after reciprocal crossing between *ire1a/b* and *ire1a/b ire1c*/+ mutants (Table 1). Transgenic plants carrying *IRE1C* promoter-driven *GUS* reporter gene construct (Fig. S2A) showed that IRE1C is expressed in anther (Fig. S3A) and embryo (Fig. S3B). No visible GUS staining was observed in vegetative tissues (root, leaf, and stem) of young seedlings with or without stress treatments (Fig. S3C). These observations are qualitatively consistent with the microarray database (Arabidopsis eFP browser; http://bar.utoronto.ca/efp/cgi-bin/efpWeb.cgi), which shows that the *IRE1C* gene is scarcely expressed in vegetative tissues. Taken together, IRE1C, which lacks a sensor domain, contributes to male gametophyte development and acts redundantly with IRE1A and IRE1B.

**Table 1.** Transmission of the *ire1c* allele through the male and female gametophytes in the progenies of the *ire1a ire1b ire1c*/+ mutants crossed with the *ire1a ire1b* mutant or self-pollinatied.

| Parental genotype |  | Genotypes of progeny |  |  |  | Observed ratio | Expected ratio |
| --- | --- | --- | --- | --- | --- | --- | --- |
| Female | Male | +/+ | c/+ | c/c | Total | +/+ : c/+ : c/c | +/+ : c/+ : c/c |
| *a/a b/b c/+* | *a/a b/b c/+* | 83 | 25 | 0 | 108 | 1.0 : 0.30 : 0<sup>a</sup> | 1 : 2 : 1 |
| *a/a b/b c/+* | *a/a b/b +/+* | 186 | 39 | 0 | 225 | 1.0 : 0.21 : 0<sup>a</sup> | 1 : 1 : 0 |
| *a/a b/b +/+* | *a/a b/b c/+* | 119 | 0 | 0 | 119 | 119 : 0 : 0<sup>a</sup> | 1 : 1 : 0 |

<sup>a</sup>Significantly different from the Mendelian segregation ratio ( $\chi^2$ , $P < 0.01$ )
+, wild-type allele; *a*, *ire1a* allele; *b*, *ire1b* allele; *c*, *ire1c* allele

**Figure 2.**
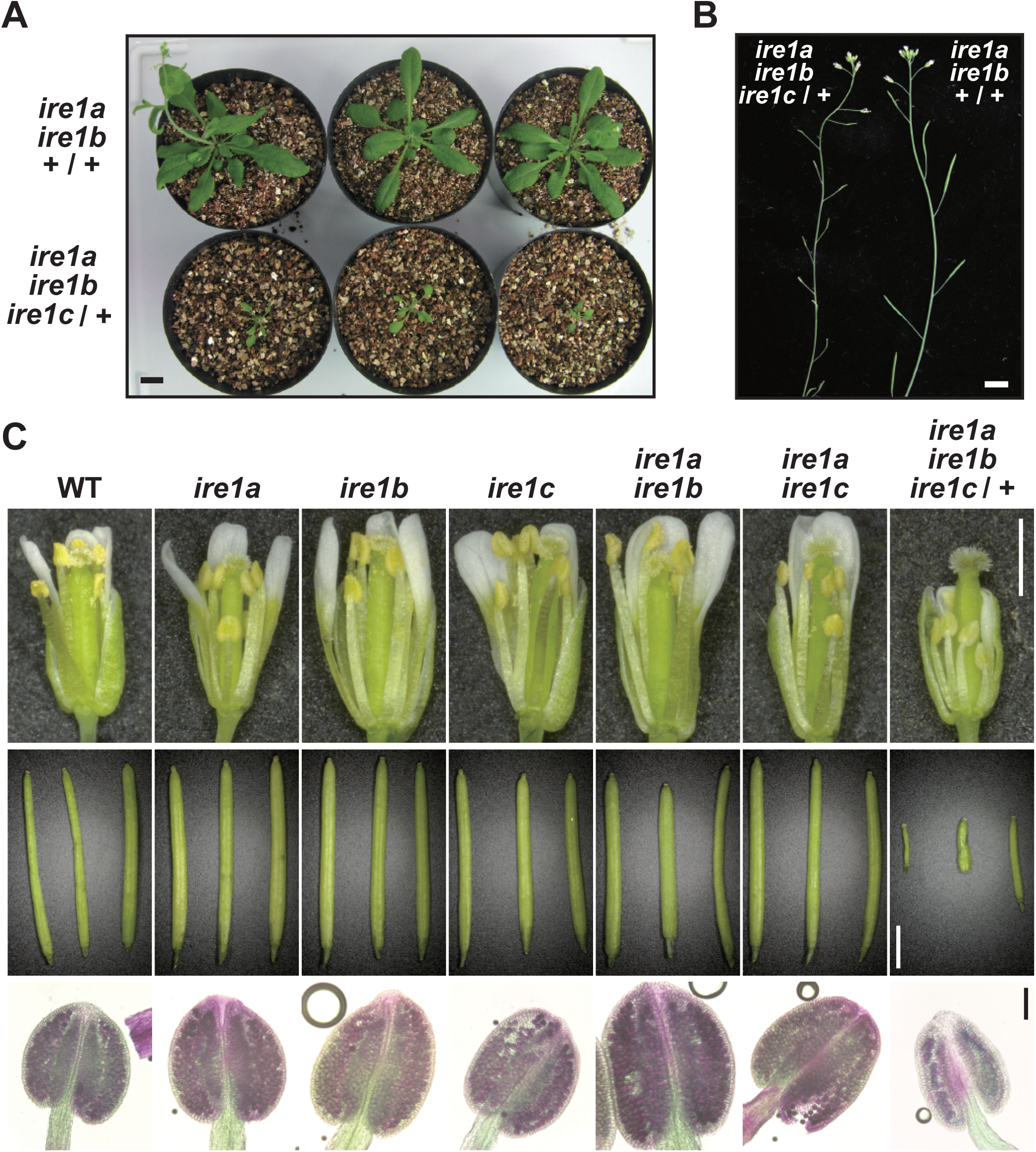
*ire1a ire1b* homozygous and *ire1c* heterozygous plants show developmental defects. (A) Self-pollinated progenies of *ire1a ire1b ire1c*/+ mutant plants at 40 DAG. Genotypes were shown on the left. Bar = 10 mm. (B) Siliques of *ire1a ire1b ire1c*/+ (left) and *ire1a ire1b* +/+ (right) plants. Bar = 10 mm. (C) Reproductive development of *ire1* mutants. Flowers at stage 14 (Smyth et al, 1990; upper; bar = 1 mm), siliques (middle; bar = 3 mm), and anthers at stage 12 stained with Alexander’s stain (lower; bar = 100 μm) from wild-type (WT) and *ire1* mutants were shown.

### Mutant IRE1B lacking the sensor domain is not responsible for UPR signal transduction

Because *ire1a/c* mutants retain fertility and IRE1-dependent UPR signal transduction, *IRE1B* possibly plays a role in both UPR and developmental processes. To investigate an unknown IRE1 function, we generated a construct expressing FLAG-tagged wild-type (WT) form of IRE1B as well as those with kinase (K487A), RNase (K821A), and luminal sensor-deletion (ΔLD) mutants under the control of its native promoter (Figs. 3A, S2B). For comparison, we generated constructs expressing FLAG-tagged wild-type (WT) IRE1A as well as mutant IRE1A with kinase (K442A) and RNase (K781A) mutation under the IRE1A native promoter (Figs. 3A, S2B). These constructs were introduced into the *ire1a/b* mutant using *Agrobacterium*-mediated transformation and T_3_ homozygous transgenic plants were used for further analyses. All of the WT and mutant IRE1 constructs expressed expected sizes of IRE1 proteins in seedlings (Fig. 3B). Up- and down-regulation of *BiP3* and *PR-4* mRNA, respectively, by Tm treatment were restored in the FLAG-IRE1A(WT) and FLAG-IRE1B(WT) transgenic plants, but not in kinase-, RNase-, and ΔLD-expressing transgenic plants (Fig. 3C). We next analyzed *in vivo* phosphorylation of FLAG-IRE1B under ER stress by Phos-tag-based western blot (Yang et al, 2010). A slower migrating, phosphorylated form of IRE1B was detected in Tm- and DTT-treated FLAG-IRE1B(WT) plants but not in the DTT-treated K487A plants (Fig. 3D). Consistent with the results of *BiP3* and *PR-4* expressions, expression of FLAG-IRE1B(WT) restored hypersensitivity of the *ire1a/b* mutant to DTT to the level observed in WT, but expression of ΔLD did not (Fig. 3E). These results indicate that the mutant IRE1B lacking the sensor domain does not contribute to the ER stress response.

**Figure 3.**
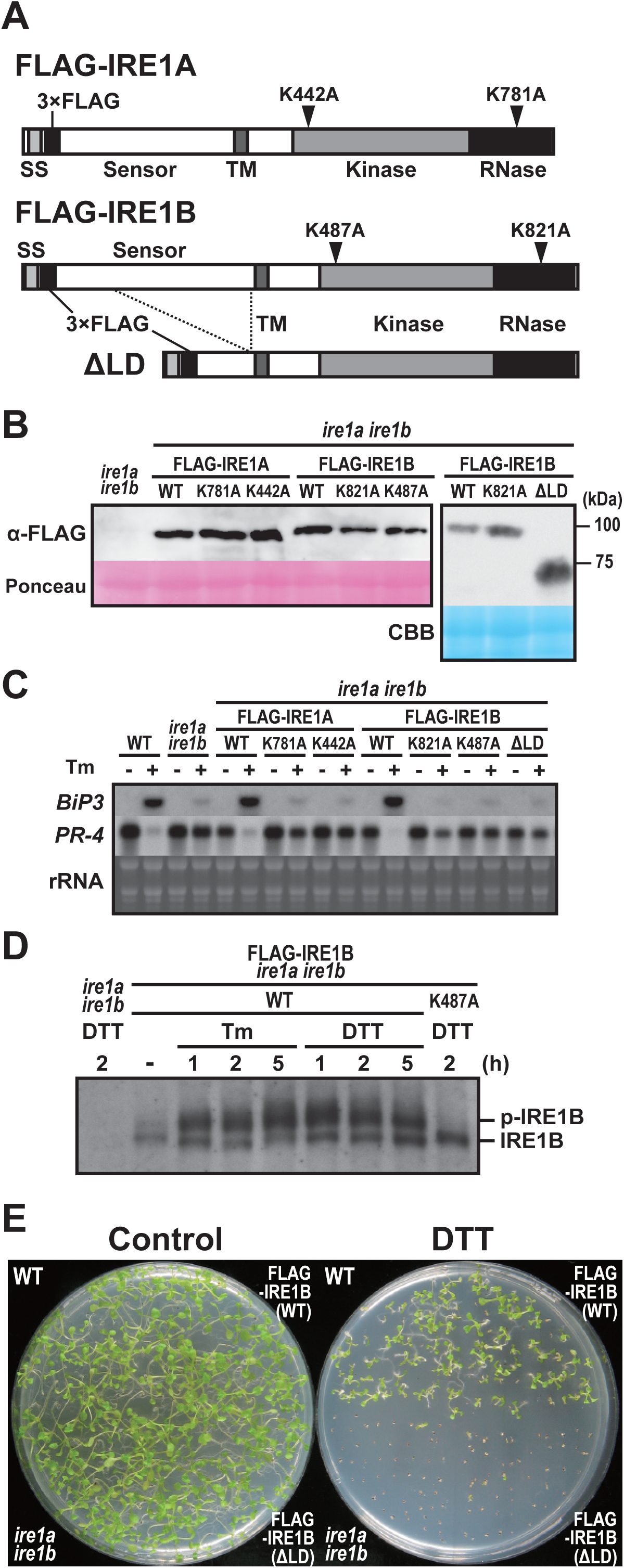
Transgenic *ire1a ire1b* plants expressing FLAG-tagged wild-type or mutant IRE1. Schema of FLAG-tagged IRE1 proteins. Mutations of kinase and RNase domains are shown as arrowheads. SS, signal sequence; TM, transmembrane domain. (B) Detection of FLAG-IRE1 proteins in the transgenic *ire1a ire1b* plants with anti-FLAG antibody. Ponceau S or CBB staining was used as loading control. (C) RNA blot analysis of *BiP3* and *PR-4* in wild-type (WT), *ire1a ire1b* mutant and transgenic *ire1a ire1b* plants. Seedlings at 10 DAG were treated with (+) or without (−) 5 mg/l tunicamycin (Tm) for 5 h. (D) Detection of FLAG-IRE1B(WT) and FLAG-IRE1B(K487A) with anti-FLAG antibody in the transgenic *ire1a ire1b* plants treated with Tm or DTT. Samples were resolved on Phos-tag SDS-PAGE to detect the phosphorylated FLAG-IRE1B (p-IRE1B). (E) DTT sensitivity of the transgenic *ire1a ire1b* plants. Seedlings at 15 DAG of the indicated lines were treated with or without 1 mM DTT.

### FLAG-IRE1B(WT) and ΔLD restore the developmental defects in *ire1a/b ire1c*/+ mutant

We crossed the *ire1a/b* mutant plants expressing FLAG-IRE1B(WT) or ΔLD with *ire1a/b ire1c*/+ mutant plants. F_1_ plants heterozygous for *IRE1C* (*ire1a/b ire1c*/+) were selected and self-pollinated. Among F_2_ plants, plants that are homozygous for the transgenes and heterozygous for *IRE1C* were selected for further analyses. Growth defects (Figs. 4A-F) and the reduction of seed set (Figs. 4G-I) in the *ire1a/b ire1c*/+ mutant were restored by expression of FLAG-IRE1B(WT) and ΔLD plants. Abortion of pollen development in *ire1a/b ire1c*/+ mutant was restored, as well (Figs. 4J-L). At the completion of meiosis, the *ire1a/b ire1c*/+ mutant plants expressing FLAG-IRE1B(WT) or ΔLD produced four viable microspores in each tetrad (Figs. 4M, N), whereas the *ire1a/b ire1c*/+ mutant frequently produced abnormal tetrads (Fig. 4O).

**Figure 4.**
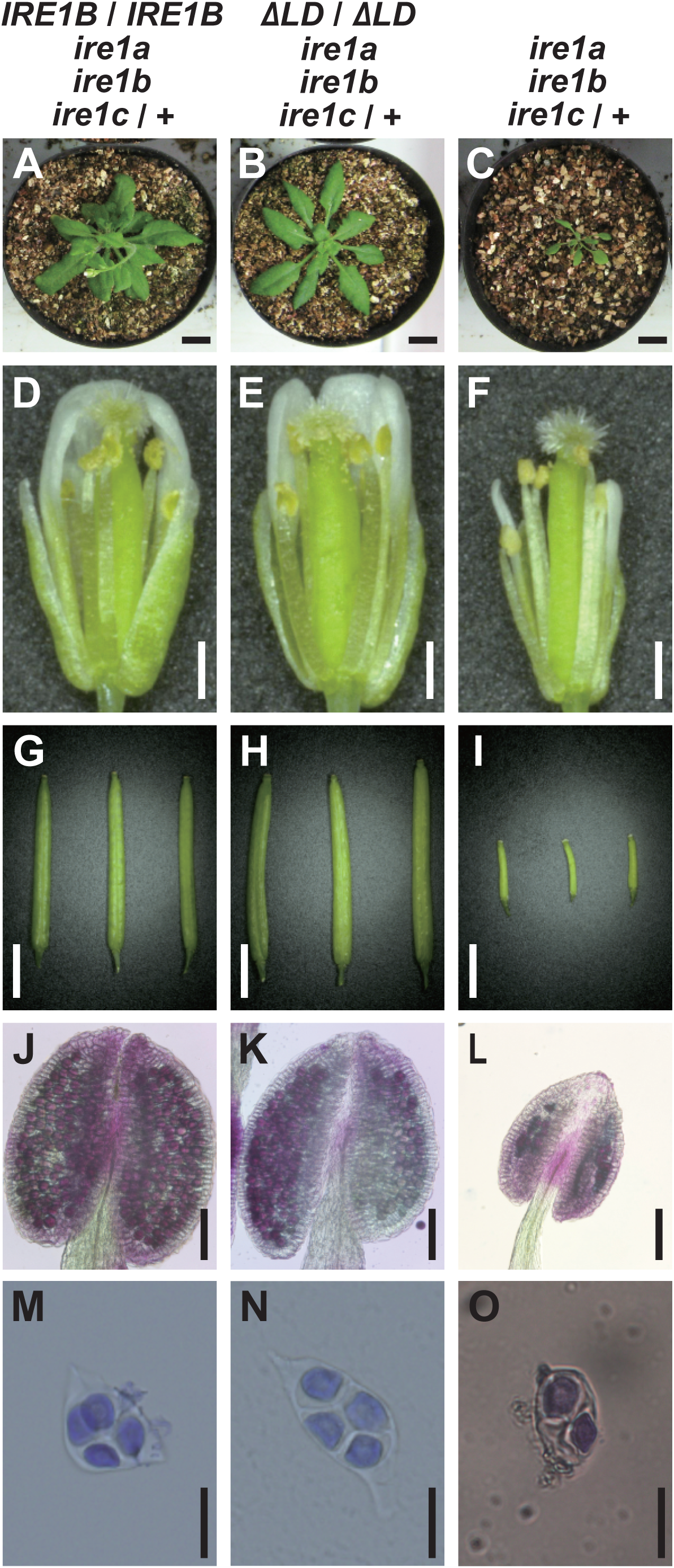
Phenotypic complementation of *ire1a ire1b ire1c*/+ mutants by FLAG-IRE1B(WT) or ΔLD. Phenotypes of the transgenic *ire1a ire1b ire1c*/+ plant having FLAG-IRE1B(WT) (left), ΔLD (center) and *ire1a ire1b ire1c*/+ plant (right). (A-C) Plants at 40 DAG. Bar = 10 mm. (D-F) Flowers at stage 14. Bar = 500 μm. (G-I) Siliques. Bar = 3 mm. (J-L) Anthers at stage 12 stained with Alexander’s stain. Bar = 100 μm. (M-O) Tetrads stained with toluidine blue. Bar = 20 μm.

### Impaired pollen development in *ire1a/b/c* gametophyte is restored by ΔLD

Unexpectedly, self-pollinated progenies of the *ire1a/b ire1c*/+ plants expressing FLAG-IRE1B(WT) or ΔLD plants segregated with the *IRE1C* allele in ratios of 1.0:1.9:0 (+/+:*c*/+:*c*/*c*; n = 154) and 1.0:1.4:0 (n = 237), respectively (Table 2). Nevertheless, their occurrence ratios of the heterozygous allele were higher than that in the *ire1a/b ire1c*/+ mutant (1.0:0.30:0; n = 108; Table 1). To determine whether the impaired transmission of the *ire1c* allele through male gametophyte in *ire1a/b ire1c*/+ mutant was restored by FLAG-IRE1B(WT) and ΔLD, we performed reciprocal crosses between the *ire1a/b ire1c*/+ plants with homozygous FLAG-IRE1B(WT) or ΔLD transgene and wild-type plants. Consistent with the results of the reciprocal crossing between *ire1a/b* and *ire1a/b ire1c*/+ (Table 1), control reciprocal crossing between wild-type and *ire1a/b ire1c*/+ showed no transmission of the *ire1c* allele through male gametophyte (Table 2). In the case of the crossing between the wild-type plants as female parents and the *ire1a/b ire1c*/+ plants with FLAG-IRE1B(WT) or ΔLD as male parents, progenies having *ire1c*/+ allele were obtained in the ratios of 1.0:0.33 (+/+:*c*/+; n = 57) and 1.0:0.57 (n = 94), respectively (Table 2). These results indicate that not only FLAG-IRE1B(WT) but also ΔLD can compensate for impaired male gametogenesis in the *ire1a/b/c* haplotype.

**Table 2.** Transmission of the *ire1c* allele through the male and female gametophytes in the progenies of the *ire1a ire1b ire1c*/+ mutants having *IRE1B* or *ΔLD* transgenes crossed with the wild-type or self-pollinated.

| Parental genotype |  | Genotypes of progeny |  |  |  | Observed ratio | Expected ratio |
| --- | --- | --- | --- | --- | --- | --- | --- |
| Female | Male | +/+ | c/+ | c/c | Total | +/+ : c/+ : c/c | +/+ : c/+ : c/c |
| +/+ +/+ +/+ | *a/a b/b c/+* | 202 | 0 | 0 | 202 | 202 : 0 : 0<sup>a</sup> | 1 : 1 : 0 |
| +/+ +/+ +/+ | *a/a b/b c/+ IRE1B* | 43 | 14 | 0 | 57 | 1.0 : 0.33 : 0<sup>a</sup> | 1 : 1 : 0 |
| +/+ +/+ +/+ | *a/a b/b c/+ $\Delta LD$* | 60 | 34 | 0 | 94 | 1.0 : 0.57 : 0<sup>a</sup> | 1 : 1 : 0 |
| *a/a b/b c/+* | +/+ +/+ +/+ | 266 | 81 | 0 | 347 | 1.0 : 0.30 : 0<sup>a</sup> | 1 : 1 : 0 |
| *a/a b/b c/+ IRE1B* | +/+ +/+ +/+ | 130 | 32 | 0 | 162 | 1.0 : 0.25 : 0<sup>a</sup> | 1 : 1 : 0 |
| *a/a b/b c/+ $\Delta LD$* | +/+ +/+ +/+ | 135 | 25 | 0 | 160 | 1.0 : 0.19 : 0<sup>a</sup> | 1 : 1 : 0 |
| *a/a b/b c/+ IRE1B* | *a/a b/b c/+ IRE1B* | 53 | 101 | 0 | 154 | 1.0 : 1.9 : 0<sup>a</sup> | 1 : 2 : 1 |
| *a/a b/b c/+ $\Delta LD$* | *a/a b/b c/+ $\Delta LD$* | 97 | 140 | 0 | 237 | 1.0 : 1.4 : 0<sup>a</sup> | 1 : 2 : 1 |

<sup>a</sup>Significantly different from the Mendelian segregation ratio ( $\chi^2$ , $P < 0.01$ )
+, wild-type allele; *a*, *ire1a* allele; *b*, *ire1b* allele; *c*, *ire1c* allele; *IRE1B*, *FLAG-IRE1B(WT)* transgene (homozygote); $\Delta LD$ , $\Delta LD$ transgene (homozygote)

To investigate the defect of male gametogenesis in the *ire1a/b/c* haplotype, we observed cross sections prepared from inflorescences of wild-type, *ire1a/b ire1c*/+, and *ire1a/b ire1c*/+ expressing ΔLD at different developmental stages. The anther size and the number of pollen grains were reduced in *ire1a/b ire1c*/+ compared to wild-type (Fig. 5). Regarding pollen development, no obvious differences were observed between *ire1a/b ire1c*/+ and wild-type at stage 8 and 9 (Fig. 5). However, a part of pollen grains collapsed in *ire1a/b ire1c*/+ at stage 11 (Fig. 5; indicated by arrowheads). The collapsing pollen grains were also shown in *ire1a/b ire1c*/+ expressing ΔLD, but the frequency was very low (Fig. 5; arrowhead). In *ire1a/b ire1c*/+ expressing ΔLD, the anther size was restored to the wild-type level (Fig. 5).

**Figure 5.**
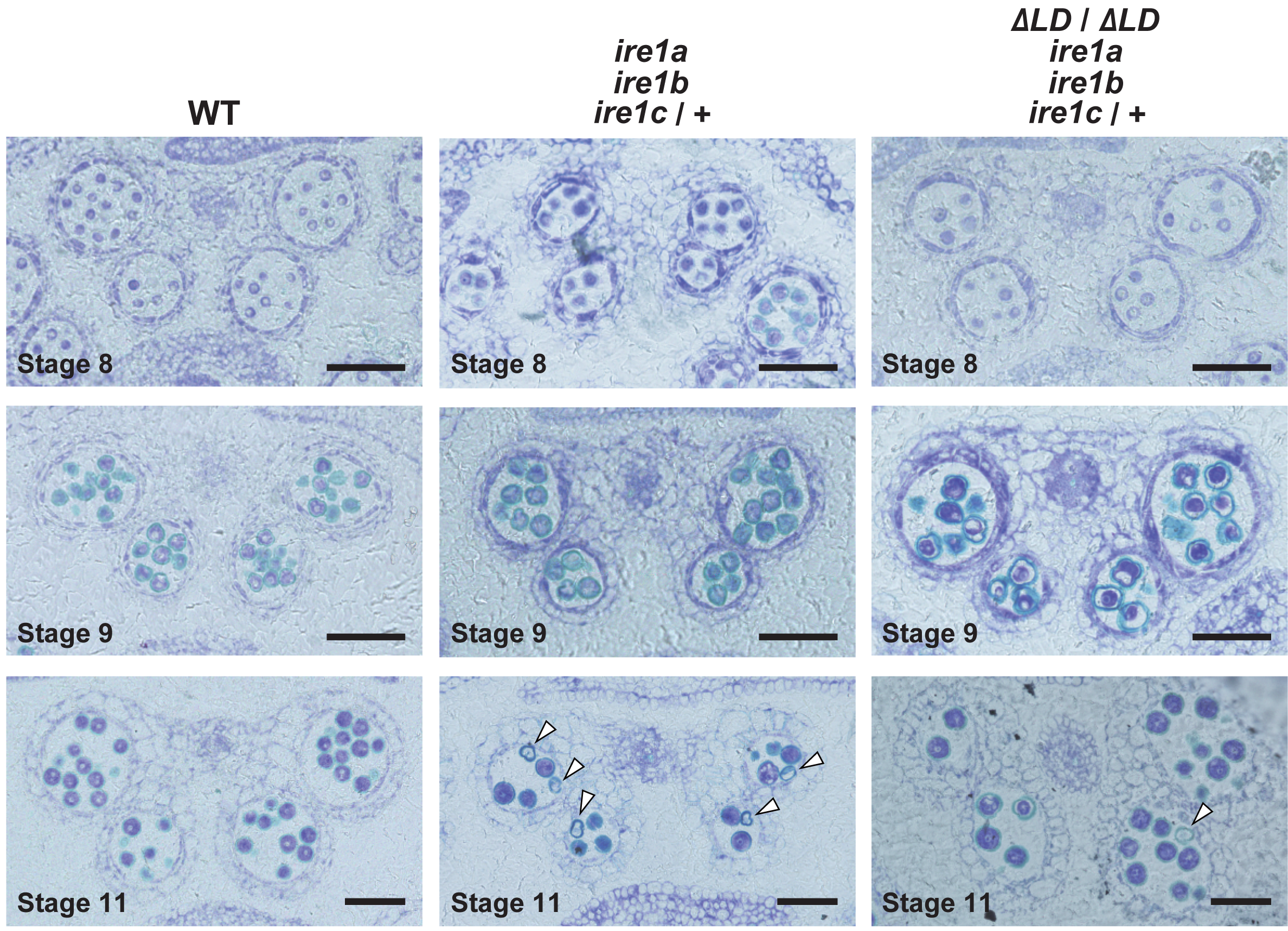
Abnormal pollen development in *ire1a ire1b ire1c*/+ is partially complemented by ΔLD. Transverse sections of developing anthers at stages 8, 9, 11 in WT (left), *ire1a ire1b ire1c*/+ mutant (middle) and transgenic *ire1a ire1b ire1c*/+ plants having ΔLD (right). Arrowheads indicate collapsed pollen grains. Bar = 50 μm.

Results of the crossing between *ire1a/b ire1c*/+ as female parents and *ire1a/b* (Table 1) or wild-type (Table 2) as male parents suggest incomplete female gametogenesis in the *ire1a/b/c* haplotype. However, *FLAG-IRE1B(WT)* and *ΔLD* transgenes did not affect the occurrence ratios of heterogeneous *ire1c*/+ allele through the female gametophyte (Table 2).

### Different IRE1 activation states by saturated fatty acids in the presence or absence of sensor domain

Growing evidence suggests that yeast and metazoan IRE1 have lipid-dependent activation machinery (Volmer & Ron, 2015). Since exogenous application of glycerol is known to reduce oleic acid (18:1) level in Arabidopsis (Kachroo *et al*, 2004), we applied glycerol treatment to Arabidopsis seedlings to increase saturated fatty acid composition. As expected, levels of palmitic acid (16:0) and stearic acid (18:0) were increased after three days of glycerol treatment in wild-type and *ire1a/b* seedlings (Fig. 6A). Glycerol treatment induced *bZIP60* splicing in wild-type but not in *ire1a/b* seedlings (Fig. 6B). Impaired *bZIP60* splicing in *ire1a/b* was restored by expression of FLAG-IRE1A(WT) and FLAG-IRE1B(WT), but not by that of kinase, RNase, and ΔLD mutants (Figs. 6C, S4A, S4B). Accumulation and phosphorylation of the FLAG-IRE1B(WT) protein was observed in the glycerol-treated seedlings (Fig. 6D). Thus, IRE1’s kinase, RNase, and sensor domains are responsible for *bZIP60* splicing under glycerol treatment *in vivo*.

**Figure 6.**
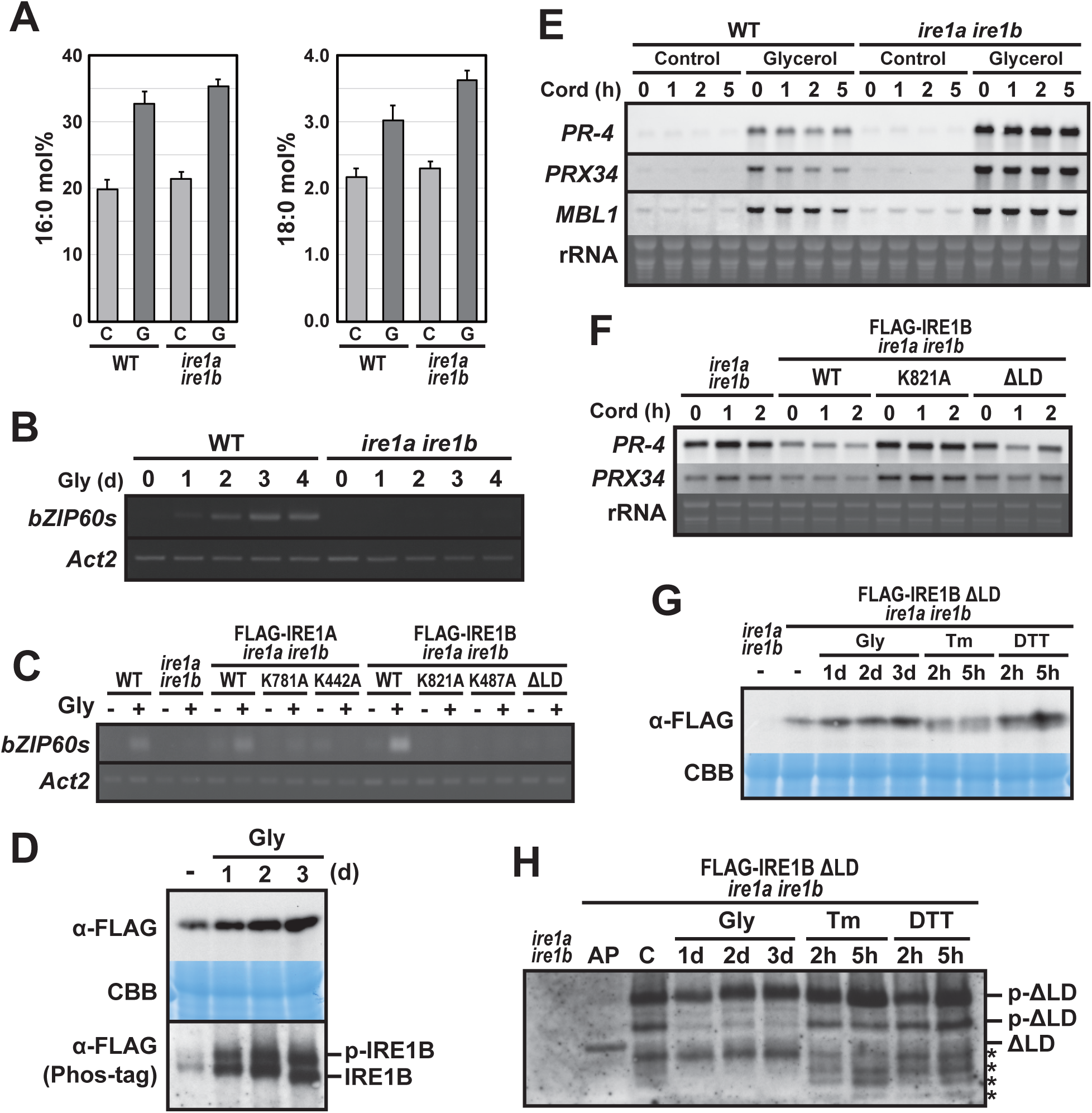
Glycerol treatment stimulates IRE1 kinase and RNase activities. (A) Percentages of saturated fatty acids (16:0 and 18:0) in WT and *ire1a ire1b* plants at 10 DAG treated with (G) or without (C) glycerol for 3 d. Error bars represent SD (n = 6). (B, C) Detection of *bZIP60* mRNA splicing in WT, *ire1a ire1b* plants (B) and FLAG-IRE1 transgenic *ire1a ire1b* plants (C) at 10 DAG. RT-PCR was performed using *bZIP60s*-specific primers. *Actin2* (*Act2*) was used as an internal control. Glycerol treatment was performed for 0-4 d. (C) Plants were treated with (+) or without (−) glycerol for 3 d. (D) Detection of FLAG-IRE1B(WT) with anti-FLAG antibody in the transgenic *ire1a ire1b* plants treated with glycerol for 0-3 d. Samples were resolved on SDS-PAGE (upper) and Phos-tag SDS-PAGE (lower) followed by immunodetection with anti-FLAG antibody. An equal loading was shown by CBB staining following SDS-PAGE (middle). (E, F) RNA blot analysis of *PR-4*, *PRX34* and/or *MBL1* in WT, *ire1a ire1b* plants (E) and FLAG-IRE1 transgenic *ire1a ire1b* plants (F) at 10 DAG. (E) Plants were treated with or without glycerol for 3 d and plants at 10 DAG were treated with cordycepin (Cord) for 0-5 h. (F) Three days of glycerol-treated plants at 10 DAG were treated with Cord for 0-2 h. (G, H) Detection of ΔLD with anti-FLAG antibody in the transgenic *ire1a ire1b* plants treated with glycerol, Tm or DTT. (G) Samples were resolved on SDS-PAGE followed by immunodetection with anti-FLAG antibody. CBB staining was used as loading control. (H) Samples were resolved on Phos-tag SDS-PAGE to detect the phosphorylated ΔLD (p-ΔLD). AP, alkaline phosphatase-treated sample. Asterisks indicate possible degradation products of ΔLD.

To determine whether glycerol treatment induces RIDD, mRNA levels of three RIDD target genes (*PR-4*, *PRX34*, and *MBL1*; Mishiba *et al*, 2013; Iwata *et al*, 2016) was analyzed in the glycerol-treated wild-type and *ire1a/b* seedlings, which were further treated with cordycepin to prevent transcription. Higher expressions of the three genes were observed in the glycerol-treated wild-type and *ire1a/b* plants compared to untreated control (Fig. 6E). Decrease in *PR-4*, *PRX34* and *MBL1* mRNA abundance was detected within 5 hours of cordycepin treatment in wild-type but not in *ire1a/b* seedlings. The impaired mRNA degradation in *ire1a/b* was restored by expression of FLAG-IRE1B(WT) and ΔLD, but not by that of RNase (K821A) mutant (Fig. 6F). Consistently, while levels of mRNA encoding cytosolic proteins, cFBPase and UGPase, did not show significant difference among the samples irrespective of glycerol treatment (Fig. S4C), expression of *PR-4*, *PRX34*, *MBL1*, and *PME41* (RIDD target; Mishiba *et al*, 2013) mRNAs was increased (P < 0.05) in *ire1a/b* and K821A compared to wild-type, FLAG-IRE1B(WT), and ΔLD plants under glycerol treatment (Fig. S4D). Accumulation of ΔLD proteins was observed under glycerol and DTT treatments (Fig. 6G). Phos-tag western blot of ΔLD protein showed two slower migrating bands than unphosphorylated protein in the untreated-, Tm- and DTT-treated plants, whereas only the slowest migrating band was detected in the glycerol-treated plants (Fig. 6H). These results indicate that glycerol treatment stimulate the mutant IRE1 proteins lacking the sensor domain, causing RIDD.

### CRISPR/Cas9-induced deletion corresponding to *IRE1B* sensor region in *ire1a/c* mutant

To gain insight into the contribution of the sensor domain-independent IRE1 activation to the developmental process, we tried to induce deletion in the sensor domain-coding region of the *IRE1B* gene in the *ire1a/c* mutant using CRISPR/Cas9 system. *Agrobacterium* harbouring pKIR1.0 binary vector (Tsutsui & Higashiyama, 2017) containing two gRNAs targeting the 5’ and 3’ ends of the IRE1B’s sensor domain-coding region (Figs. 7A, S2C) was used to transform the *ire1a/c* mutant. We selected T_2_ lines showing 3:1 segregation ratio for the presence and absence of RFP fluorescence (see Fig. S2C) in seeds and picked up some seeds with no fluorescence, indicative of no T-DNA insertion for further analyses (Tsutsui & Higashiyama, 2017). When we amplify an *IRE1B*-coding genomic region by PCR, we obtained smaller bands than that would be expected from intact *IRE1B* in some T_2_ plants, indicating that deletion is successfully introduced. Therefore, their self-progenies (T_3_) were used for further analyses. In lines #2-5 and #9-6, all T_3_ plants analyzed showed a single, smaller *IRE1B* fragment, indicating homozygous deletion of *IRE1B* locus (Fig. S5A). Sequence analysis showed deletion of 981 and 1,216 bp regions, each corresponding to part of the sensor domain in #2-5 and #9-6, respectively, and 1 bp deletion at the gRNA2 target site was also detected in #2-5 (Fig. 7A). Consistently, RT-PCR showed expression of shorter *IRE1B* mRNA in #2-5 and #9-6 seedlings (Fig. S5B). We expected that two isolated lines express N-terminally truncated IRE1B proteins by illegitimate translation (Makino *et al*, 2016), because translation from the original ATG produces premature N-terminal peptides (52 AA and 76 AA, respectively). These predicted ORFs in #2-5 and #9-6 have a truncated and an intact transmembrane domain, respectively (Fig. 7A). High sensitivity to Tm and DTT equivalent to *ire1a/b* were found in #2-5 and #9-6 lines compared to that in *ire1a/c* (Fig. 7B, S5C). Like the *ire1a/b* mutant, up- and down-regulation of *BiP3* and *PR-4* mRNA, respectively, by Tm treatment were diminished in #2-5 and #9-6 lines (Fig. 7C). Glycerol treatment-dependent *bZIP60* splicing as shown in *ire1a/c* was also diminished in #2-5 and #9-6 lines (Fig. 7D). Nevertheless, mRNA expression of RIDD target genes in these lines was decreased as compared to those in *ire1a/b* under glycerol treatment (Fig. 7E, S6A). Growth defects and the reduction of seed set, which occurred in *ire1a/b ire1c*/+ mutant, were not observed in the #2-5 and #9-6 plants (Fig. S6B, S6C).

**Figure 7.**
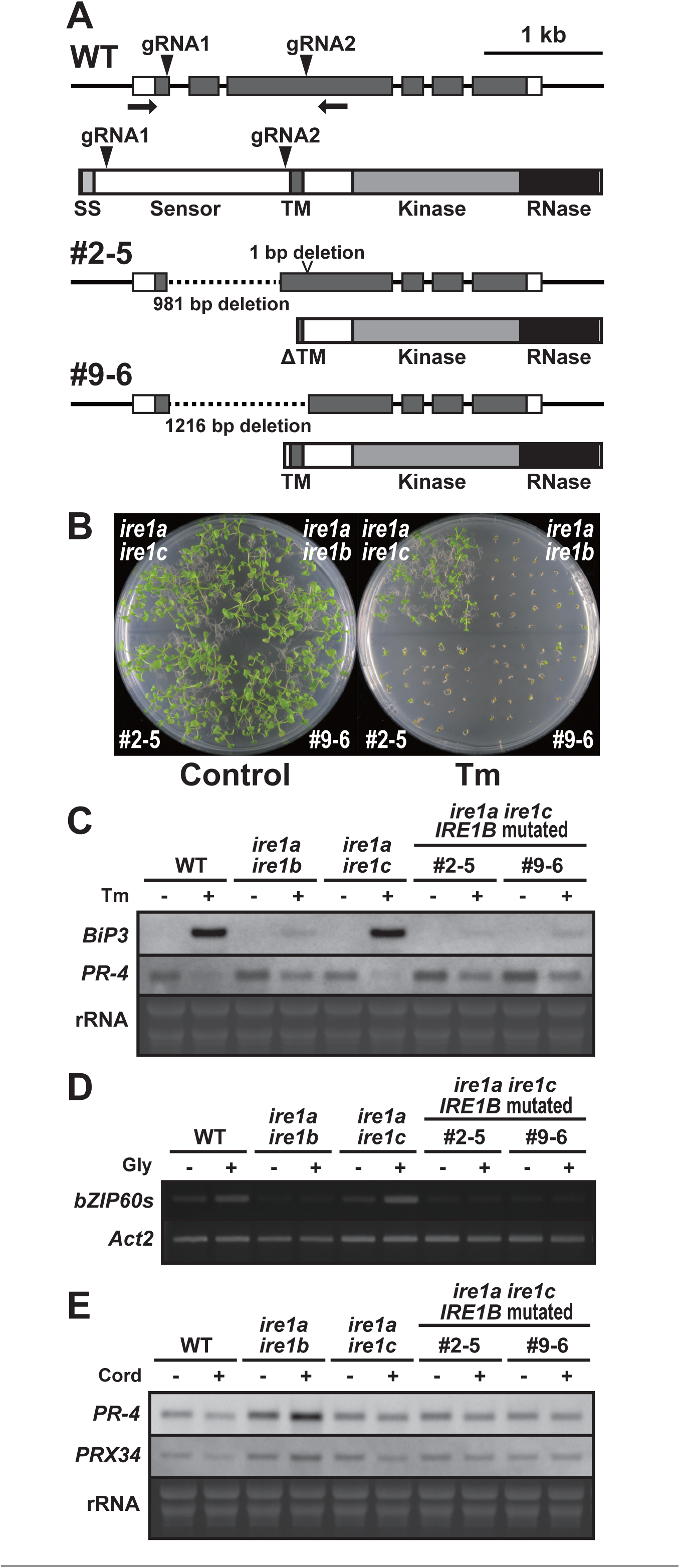
CRISPR/Cas9-mediated *IREB* gene editing in *ire1a ire1c* mutant. (A) Schema of hypothetical IRE1B products in the *IRE1B*-mutated *ire1a ire1c* T_3_ plant lines #2-5 and #1-10. Positions of gRNA target sites are shown as arrowheads. SS, signal sequence; TM, transmembrane domain. (B) Tm sensitivity of the *ire1a ire1b*, *ire1a ire1c*, and *IRE1B*-mutated *ire1a ire1c* plants. Seedlings at 15 DAG of the indicated lines were treated with or without 0.1 mg/l Tm. (C) RNA blot analysis of *BiP3* and *PR-4* in WT, *ire1a ire1b*, *ire1a ire1c* mutant and *IRE1B*-mutated *ire1a ire1c* plants. Seedlings at 10 DAG were treated with (+) or without (−) 5 mg/l tunicamycin (Tm) for 5 h. (D) Detection of *bZIP60* mRNA splicing in WT, *ire1a ire1b*, *ire1a ire1c*, and *IRE1B*-mutated *ire1a ire1c* plants at 10 DAG treated with (+) or without (−) glycerol for 3 d. (E) RNA blot analysis of *PR-4* and *PRX34* in WT, *ire1a ire1b*, *ire1a ire1c*, and *IRE1B*-mutated *ire1a ire1c* plants at 10 DAG treated with glycerol for 3 d. The samples were treated with (+) or without (−) Cord for 2 h immediately before sampling.

## Discussion

IRE1 is known as the most conserved and sole UPR signal transducer in lower eukaryotes (Mori, 2009). Evolution of multicellular organisms adapt IRE1 functions not only to environmental conditions but also to developmental conditions, as in the fact that IRE1 deficiency causes embryonic lethality in some organisms. In developmental processes, specific cells producing a large amount of secretory proteins, such as, β-cells of pancreas (Lee *et al*, 2011), goblet cells (Tsuru *et al*, 2013), and dendritic cells (Osorio *et al*, 2014), activate IRE1 in normal condition. These findings raise a question of whether production of unfolded proteins is prerequisite for the IRE1 activation in these specific cells. The present study showed that IRE1 activation without sensing unfolded protein is required for multifaceted developmental processes in Arabidopsis. We speculate that unfolded protein-independent IRE1 activation is a feature of anticipatory UPR (Vitale & Boston, 2008; Rutkowski & Hegde, 2010) to avoid producing ‘unprofitable’ unfolded proteins, as a primer for UPR, during the evolution of an unfolded protein-sensing system in multicellular organisms.

We found that plants heterozygous for the *IRE1C* allele (*ire1c*/+) in *ire1a/b* mutant background display developmental defects of male (and also probably female) gametogenesis, incomplete floral organ formation, and retardation of vegetative growth (Fig. 2). The incomplete dominance of the *IRE1C* allele is probably due to low expression of *IRE1C* transcript. The *IRE1C* expression is distinct in anther (Fig. S3A) and embryo (Fig. S3B), which is somewhat similar to that of *IRE1B* (Koizumi *et al*, 2001). Expression of *IRE1A* alone may be insufficient for proper Arabidopsis development because we could not obtain *ire1b ire1c* double mutant. Functions and physiological significance of the Arabidopsis IRE1C is still unclear. A closely related species, *Arabidopsis lyrata*, has four *IRE1* genes, *IRE1A* (XP_002884063), *IRE1B* (EFH48369), *IRE1C* (XP_002884871), and *IRE1C-like* (EFH64123). Amino acid sequence similarities between those in *A. thaliana* and *A. lyrata* are high in IRE1A (93%) and IRE1B (91%), whereas IRE1C proteins are more diversified (66%). *A. lyrata* also has *IRE1C-like* gene with low sequence similarity. These results suggest that these *IRE1C* genes probably arose through gene duplication during evolution of Brassicaceae species.

Pollen is known to be particularly sensitive to environmental conditions that disturb protein homeostasis (Fragkostefanakis *et al*, 2016). In high temperature, *ire1a/b* mutant displays male-sterility (Deng *et al*, 2016), suggesting that pollen development is sensitive to heat stress and that IRE1-dependent UPR pathway is required for protecting male fertility from heat stress. This feature seems to be distinct from the requirement of the UPR-independent IRE1 activation for pollen development in unstressed condition observed in the present study. IRE1 activation in pollen without stress conditions was suggested by detection of bZIP60s in anther (Iwata *et al*, 2008), while *ire1a/b* mutant does not compromise pollen development under normal conditions (Deng *et al*, 2013, 2016). In the present study, we demonstrate that IRE1C, which lacks a sensor domain, acts redundantly with IRE1A and IRE1B in pollen development. Observation of the pollen development in *ire1a/b ire1c*/+ mutant showed reduced number of pollen (Fig. 5) and abnormal tetrad (Fig. 4O), suggesting that male gametogenesis in the *ire1a/b/c* haplotype is defective in meiosis. Additionally, the *ire1a/b ire1c*/+ mutant showed collapsed pollen grains at stage 11 (Fig. 5), which is somewhat similar to that observed in RNAi-mediated suppression of ER- and Golgi-located phospholipase A_2_ transgenic plants (Kim *et al*, 2011). Together with the results that no transmission of *ire1c* allele were found through *ire1a/b* male gametophyte and that ΔLD can restore the transmission in the *ire1a/b/c* haplotype (Tables 1, 2), the unfolded protein-independent IRE1 activation is required for the male gametogenesis in unstressed conditions.

Genetic analysis also showed distorted segregation ratios in progenies of crosses between *ire1a/b ire1c*/+ females and *ire1a/b* or wild-type males (Tables 1, 2), suggesting that the unfolded protein-independent IRE1 activation may be involved in embryogenesis or female gametogenesis. This hypothesis is supported by the observations that the sensor domain-lacking *IRE1B* mutant lines (i.e., #2-5 and #9-6) in *ire1a/c* background set seeds normally (Fig. S6C), and that *IRE1B* (Koizumi *et al*, 2001) and *IRE1C* (Fig. S3B) express in ovule and embryo, respectively. Inconsistent with the result of the #2-5 and #9-6 lines, we could not obtain homozygous *ire1c* mutant plants from selfed progenies of the *ire1a/b ire1c*/+ plants expressing FLAG-IRE1B(WT) or ΔLD plants. A possible reconciliation could be that the *IRE1B* transgene promoter is not expressed in embryo at a level equivalent to endogenous *IRE1B* due to epigenetic modifications of the promoter or to insufficiency in the length of the transgene promoter.

By co-expression of two gRNAs and Cas9, part of *IRE1B*-coding regions (981 and 1,216 bp) corresponding to its sensor domain was removed from the *ire1a/c* mutant genome (Fig. 7A). It is inconceivable that illegitimate translation that results in sensor domain-lacking IRE1B (Fig. 7A) does not occur, because the *ire1a/b/c* mutant is lethal. *IRE1B* mRNA may be compatible with illegitimate translation because its 5’UTR contains uORF and *IRE1B* mRNA degradation by premature stop codon (Garneau *et al*, 2007) was not observed in #2-5 and #9-6 (Fig. S5B). Therefore, it is most conceivable that expression of sensor domain-lacking IRE1B confers normal seed set in #2-5 and #9-6. This unexpected IRE1 translation may also occur in known *ire1* null mutants in other organisms, such as *C. elegans ire1(v33)* mutant (Shen *et al*, 2001). Whether a shorter TM domain in #2-5 is functional at a transmembrane domain needs to be elucidated.

The present study shows distinct modes of IRE1 activation by saturated fatty acid *in vivo*. Compared to Tm or DTT treatments (Mishiba *et al*, 2013), the level of IRE1 activation (i.e. fold induction of *bZIP60s* mRNA) by glycerol treatment was low. While full-length IRE1B activates both *bZIP60* splicing and RIDD under glycerol treatment, sensor domain-lacking IRE1B activates RIDD but not *bZIP60* splicing (Figs. 6, 7, S4, and S6). Biochemical studies in other model systems showed that oligomerization of IRE1 is required for XBP1/HAC1 cleavage but not RIDD (Tam *et al*, 2014). If plant IRE1 also acts in the same way, sensor domain-lacking IRE1 may be less likely to undergo oligomerization by saturated fatty acid. *In vivo* phosphorylation of FLAG-IRE1B was found under glycerol treatment (Fig. 6D), whereas phosphorylation of ΔLD is rather complicated. Since ΔLD exhibits multiple phosphorylation states and one of the state is stable regardless of stress (Fig. 6H), basal RIDD activity (Maurel *et al*, 2014) may exist in unstressed tissues. This hypothesis may explain growth retardation of *ire1a/b ire1c*/+ plants (Fig. 2A) in unstressed conditions. Considering our current observations that ΔLD restores developmental defects found in the *ire1a/b ire1c*/+ mutant and that deletion of the IRE1B’s sensor domain in #2-5 and #9-6 does not prevent their development, RIDD activity may play an important role in the developmental processes. This hypothesis is partially supported by the findings of Deng *et al*, (2013) that *ire1a/b bzip28* but not *bzip60 bzip28* mutant haplotypes impaired male gametogenesis (note that bZIP28 is another UPR arm in Arabidopsis).

In conclusion, this study shows that the unfolded protein-independent IRE1 activation is involved in multifaceted developmental processes, especially pollen development, in Arabidopsis. We hypothesize that the alternative IRE1 activation pathway may be conserved in multicellular organisms as an ‘anticipatory’ mode (Walter & Ron, 2011; Shapiro *et al*, 2016) of the UPR to avoid producing unfolded proteins in differentiated cells synthesizing a large amount of secretory proteins.

## Materials and Methods

### Plant materials and stress treatments

*Arabidopsis thaliana* Col-0 ecotype and T-DNA insertion mutants in the Col-0 background were used in this study. Plants were grown on soil or half strength Murashige and Skoog (1/2 MS) medium containing 0.8% agar and 1% sucrose under 16 h light and 8 h dark conditions at 22°C. T-DNA insertion mutants of *ire1a*, *ire1b*, and *ire1a/b* were described previously (Nagashima *et al*, 2011). A T-DNA insertion mutant of *IRE1C* (SALK_204405) was obtained from the Arabidopsis Biological Resource Center. T-DNA insertions were confirmed by genomic PCR as shown in Fig. S1A and S1B using primers listed in Table S1. Extraction of DNA for genotyping was carried out as described by Kasajima *et al*. (2004). Genotyping PCR was performed using KAPA Taq Extra PCR kit (Kapa Biosystems) according to the manufacturer’s protocol. To test the sensitivity of seedlings to Tm and DTT, sterilized seeds were sown on a 1/2 MS plate containing Tm (0.1 μg/mL) or DTT (1 mM). For stress treatments, 10-d-old seedlings in 1/2 MS liquid medium (Nagashima *et al*, 2011) were treated with 5 μg/mL Tm, 1 mM DTT, or DMSO (mock) for 1 to 5 h. For glycerol treatment, 7-d-old seedlings in liquid medium were treated with 50 mM glycerol for 3 d followed by treatment with 0.6 mM cordycepin (Wako) for 0 to 5 h.

### Production of transgenic Arabidopsis plants

For *IRE1C* promoter-*GUS* fusion construct, a 1,577-bp fragment of the IRE1C promoter was cloned into pENTR/D-TOPO (Thermo Fisher Scientific) and transferred into pSMAB-GW-GUS (Fig. S2A) binary vector by Gateway LR reaction (Thermo Fisher Scientific). For FLAG-tagged IRE1 constructs, 5,126 and 5,081 bp fragments of *IRE1A* and *IRE1B* genes, respectively, comprising approximately 0.9 and 1.2 kb of their promoter regions, respectively, were amplified by PCR with primers listed in Table S1 and cloned into pENTR/D-TOPO. Triple FLAG-tag and mutations in the kinase and RNase domains were introduced by PCR. These pENTR vectors were transferred into pSMAB-GW destination binary vector (Fig. S2B) by Gateway LR reaction. The ΔLD-expressing vector was made by partial *Mlu*I digestion of pSMAB-FLAG-IRE1B(WT) followed by *Nhe*I digestion, blunt end formation by T4 DNA polymerase, and self-ligation. For CRISPR/Cas9, we used pKIR1.0 binary vector (Tsutsui & Higashiyama, 2017) comprising AtU6 promoter-driven gRNA1 and AtU6 promoter-driven gRNA2 (Fig. S2C). The target sequences of the gRNA1 and gRNA2 are listed in Table S1. The binary vectors were introduced into *Rhizobium* strain EHA101 (Hood *et al*, 1986) and transformed into *ire1a/b* (for *IRE1A* and *IRE1B* constructs), *ire1a/c* (for CRISPR/Cas9) mutants, and wild-type (for *IRE1C* promoter-*GUS*) by the floral dip method (Clough & Bent, 1998).

### RNA analysis

Total RNA was extracted using a NucleoSpin RNA kit (Takara) according to the manufacturer’s protocol. For RT-PCR and qPCR, 500 ng of RNA was subjected to reverse transcription with random primers using High Capacity cDNA Reverse Transcription Kit (Thermo Fisher Scientific) according to the manufacturer’s protocol. qPCR was performed with an ABI 7300 Real-Time PCR System (Applied Biosystems) using Thunderbird SYBR qPCR Mix (Toyobo), and the transcript abundance of the target genes were normalized to that of 18S rRNA (Zoschke *et al*, 2007). Primers used for RT-PCR and qPCR are listed in Table S1. RNA gel blot analysis was conducted using DIG High Prime DNA Labeling and Detection Starter Kit II (Roche) according to the manufacturer’s protocol. Primers used to generate probes are listed in Table S1.

### Protein analysis

Total protein extraction from Arabidopsis seedlings was performed as described by Liu *et al* (2011). Protein extracts were fractionated by SDS-PAGE followed by western blotting with HRP-conjugated anti-FLAG antibody (PM020-7; MBL; 1:10,000) and chemiluminescent detection using Chemi-Lumi One Ultra (Nacalai Tesque). To reveal Rubisco large subunit, Coomassie Brilliant Blue (CBB) staining of the gel or Ponceau-S staining of the membrane were performed. For Phos-tag SDS-PAGE (Kinoshita & Kinoshita-Kikuta, 2011), 6% polyacrylamide gels containing 5-15 μM Phos-tag acrylamide (Wako) and 10-30 μM ZnCl_2_ were run according to the manufacturer’s protocol. For Phos-tag SDS-PAGE sample preparation, Arabidopsis seedlings was ground in liquid nitrogen and homogenized in an extraction buffer (100 mM Tris-HCl, pH 7.5, 0.25 M sucrose, 5 mM PMSF). The homogenate was centrifuged at 2,000 ×g for 2 min (4°C) and the supernatant was centrifuged at 10,000 ×g for 2 min (4°C). The supernatant was further centrifuged at 100,000 ×g for 30 min (4°C). The crude microsomal fraction pellet was subjected to the protein extraction as described above.

### Histological analysis

Anther samples were fixed with 4% glutaraldehyde in 60 mM HEPES (pH 7.0) containing 0.125 M sucrose. After dehydration in a graded series of ethanol/water mixtures, the samples were embedded in Quetol 651 resin (Nisshin EM) with formulation for plant material (Ellis, 2016). Semi-thin (2 μm) transverse sections were prepared from at least eight resin blocks per sample and stained with 0.2% toluidine blue. Stained sections were examined using a BZ-9000 microscope (Keyence). Histochemical GUS staining of *IRE1C* promoter-*GUS* plants was performed as previously described (Iwata *et al*, 2008).

### Fatty acid analysis

Arabidopsis seedlings (applox. 150 mg FW) at 10 DAG was used for the analysis of fatty acid composition. The fatty acids extracted with hexane were methylated and purified with fatty acid methylation kit (Nacalai Tesque) following the manufacturer’s instructions. The fatty acid compositions were determined using an Agilent 6890 gas chromatograph (Agilent Technologies) equipped with a DB-23 column (30 m × 0.25 mm × 0.25 μm; Agilent Technologies). Nonadecanoic methyl ester (C19:0) was used as the internal standard.

## Acknowledgements

We thank Ms. Ayumi Sakei, Ms. Fumika Yagi, Ms. Sae Saito (Osaka Prefecture University), and Dr. Yukihiro Nagashima (Texas A&M University) for technical assistance. We also thank the ABRC and GABI-Kat for providing T-DNA insertion lines, Dr. Hiroki Tsutsui and Dr. Tetsuya Higashiyama (Nagoya University) for pKIR1.0. This work was supported by Grant-in-Aid for Scientific Research (26450010, 17K07610) from The Ministry of Education, Culture, Sports, Science and Technology (MEXT), Shorai Foundation for Science and Technology, and Takeda Science Foundation.

## Author contributions

KiM directed the project. KiM and NK design the experiments. KiM, YI, RH and NN produced transgenic plants. TM and KiM performed histological analysis. AM performed fatty acid analysis. KiM performed all other experiments. KiM and YI provided the materials and reagents. KiM, YI, and NK wrote the manuscript. All authors have read and approved the manuscript.

## Figure Legends for Supplementary Materials

**Figure S1. Genotyping of *IRE1* genes in *ire1* mutants.** (A) Schematic representation of T-DNA insertion sites in *ire1a*, *ire1b*, and *ire1c*. Grey and white boxes indicate coding sequences and untranslated regions, respectively. Arrows indicate the positions of primers (Table S1) used for genotyping. (B) Genotyping of wild-type, *ire1a*, *ire1b*, *ire1c*, *ire1a ire1b*, *ire1a ire1c*, and selfed siblings (*ire1c*/+ or +/+) of *ire1a ire1b ire1c*/+. Lanes 1-3, three independent plants for each mutant. Lane M, 100 bp DNA ladder. M, mutant. W, wild-type.

**Figure S2. T-DNA constructs of the binary vectors used in this study.** (A) *IRE1C* promoter-driven *GUS* reporter gene construct. *IRE1C* promoter region within pENTR vector was transferred into pSMAB-GW-GUS binary vector through Gateway LR reaction. (B) FLAG-tagged IRE1A and IRE1B constructs. Genomic regions of the IRE1A and IRE1B genes were cloned into pENTR vector and transferred into pSMAB-GW binary vector. Modifications are indicated in each panel. (C) A CRISPR/Cas9 binary vector containing gRNA1 and gRNA2 targeting the sensor domain of the *IRE1B* gene (see Fig. 7A).

**Figure S3. Tissue-specific expression of *IRE1C* gene.** (A-C) GUS histochemical staining of transgenic Arabidopsis containing *IRE1C* promoter-*GUS* fusion construct in floral tissues (A), ovules and embryo (B). Bar = 100 μm. (C) 8-d-old seedlings treated with or without tunicamycin (Tm) for 5 h, and 11-d-old seedlings treated with glycerol for 3d. Bar = 5 mm.

**Figure S4. Effect of RNase and sensor domain of IRE1B on *bZIP60* splicing and RIDD under glycerol treatment.** (A) Detection of *bZIP60s* mRNA splicing in WT, *ire1a ire1b*, and FLAG-IRE1B(WT, ΔLD) transgenic *ire1a ire1b* plants at 10 DAG. RT-PCR was performed using *bZIP60s*-specific primers. *Actin2* (*Act2*) was used as an internal control. Glycerol treatment was performed for 0, 1, and 3 d. (B-D) The relative mRNA levels of *bZIP60s* (B), cytosolic marker protein genes (C), and RIDD target genes (D) in WT, *ire1a ire1b*, and FLAG-IRE1B(WT, K821A, ΔLD) transgenic *ire1a ire1b* plants. RNA from seedlings at 10 DAG treated with (+) or without (−) glycerol for 3 d was subjected to qPCR. Data are means ± SEM of four independent experiments. Different letters within each treatment indicate significant differences (P < 0.05) by the Tukey–Kramer HSD test.

**Figure S5. Characteristics of the CRISPR/Cas9-mediated *IRE1B* mutant lines #2-5 and #9-6.** (A) Genotyping of *IRE1A-C* genes in wild-type, *ire1a ire1b*, *ire1a ire1c* mutants, and T_3_ plants of lines #2-5 and #9-6. Lane M, 100 bp DNA ladder. Lanes 1-6, six independent plants for each line. M, mutant. W, wild-type. Note that PCR amplification of *Cas9* gene was performed to detect T-DNA, and that #9-6 lane 7 is a T_2_ sibling plant having T-DNA used as a control. (B) RT-PCR of *IRE1B* mRNA in *ire1a ire1c*, *ire1a ire1b* mutants, and T_3_ plants of #2-5 and #9-6. *Actin2* (*Act2*) was used as an internal control. (C) DTT sensitivity of the *ire1a ire1b*, *ire1a ire1c* mutants, and T_3_ plants of #2-5 and #9-6. Seedlings at 15 DAG were treated with or without 1mM DTT.

**Figure S6. #2-5 and #9-6 lines retain RIDD activity and show no phenotypic abnormality.** (A) The relative mRNA levels of RIDD target genes in WT, *ire1a ire1b*, *ire1a ire1c*, and *ire1a ire1c* with *IRE1B* mutation T_3_ lines #2-5 and #9-6. RNA from seedlings at 10 DAG treated with glycerol for 3 d was subjected to qPCR. Data are means ± SEM of four independent experiments. Different letters indicate significant differences (P < 0.05) by the Tukey–Kramer HSD test. (B) T_3_ plants of lines #2-5 and #9-6 and *ire1a ire1b ire1c*/+ mutant at 40 DAG. Bar = 10 mm. (C) Siliques of the lines #2-5 (left) and #9-6 (right) plants. Bar = 3 mm.

