## Supplemental figures for "Unfolded protein-independent IRE1 activation contributes to multifaceted developmental processes in Arabidopsis"

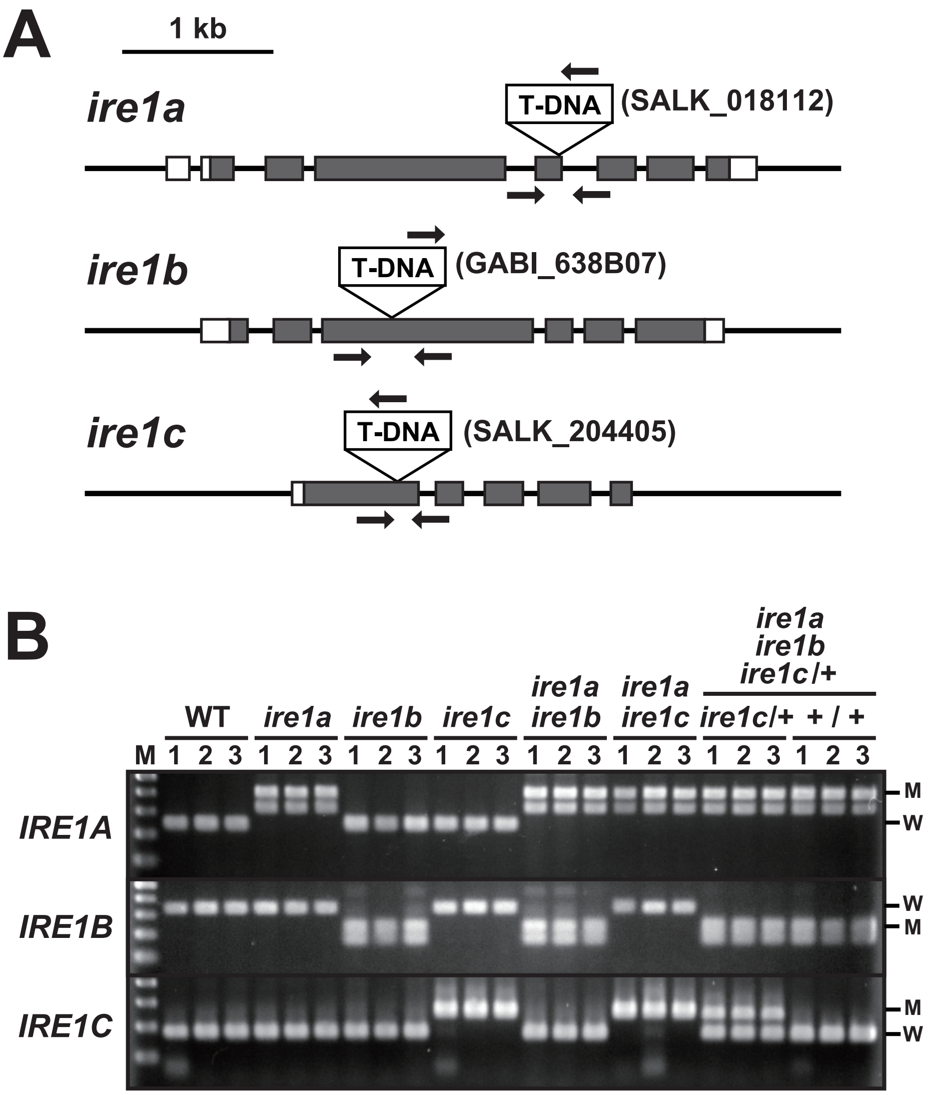


**Figure S1. Genotyping of *IRE1* genes in *ire1* mutants.** (A) Schematic representation of T-DNA insertion sites in *ire1a*, *ire1b*, and *ire1c*. Grey and white boxes indicate coding sequences and untranslated regions, respectively. Arrows indicate the positions of primers (Table S1) used for genotyping. (B) Genotyping of wild-type, *ire1a*, *ire1b*, *ire1c*, *ire1a ire1b*, *ire1a ire1c*, and selfed siblings (*ire1c*/+ or +/+) of *ire1a ire1b ire1c*/+. Lanes 1-3, three independent plants for each mutant. Lane M, 100 bp DNA ladder. M, mutant. W, wild-type.

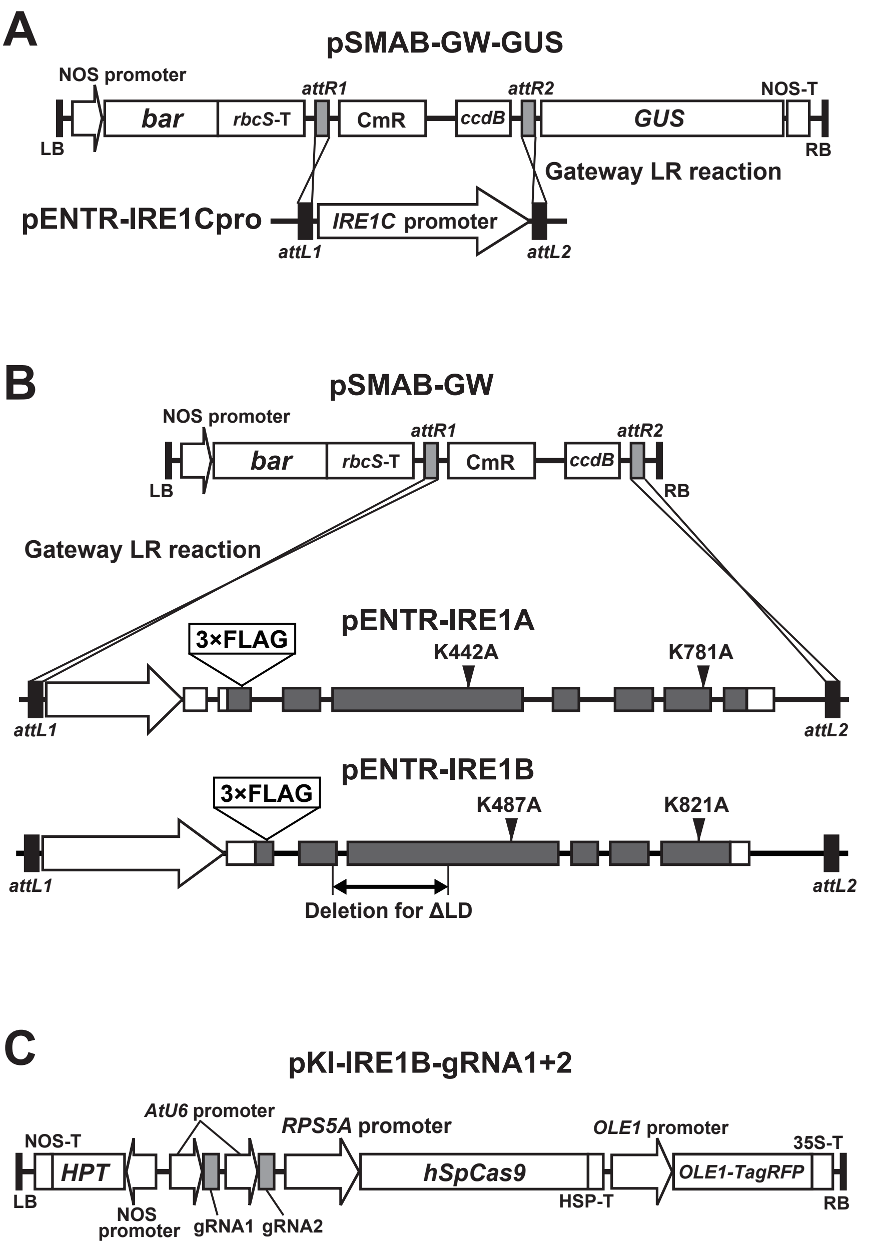


**Figure S2. T-DNA constructs of the binary vectors used in this study.** (A) *IRE1C* promoter-driven *GUS* reporter gene construct. *IRE1C* promoter region within pENTR vector was transferred into pSMAB-GW-GUS binary vector through Gateway LR reaction. (B) FLAG-tagged IRE1A and IRE1B constructs. Genomic regions of the IRE1A and IRE1B genes were cloned into pENTR vector and transferred into pSMAB-GW binary vector. Modifications are indicated in each panel. (C) A CRISPR/Cas9 binary vector containing gRNA1 and gRNA2 targeting the sensor domain of the *IRE1B* gene (see Fig. 7A).

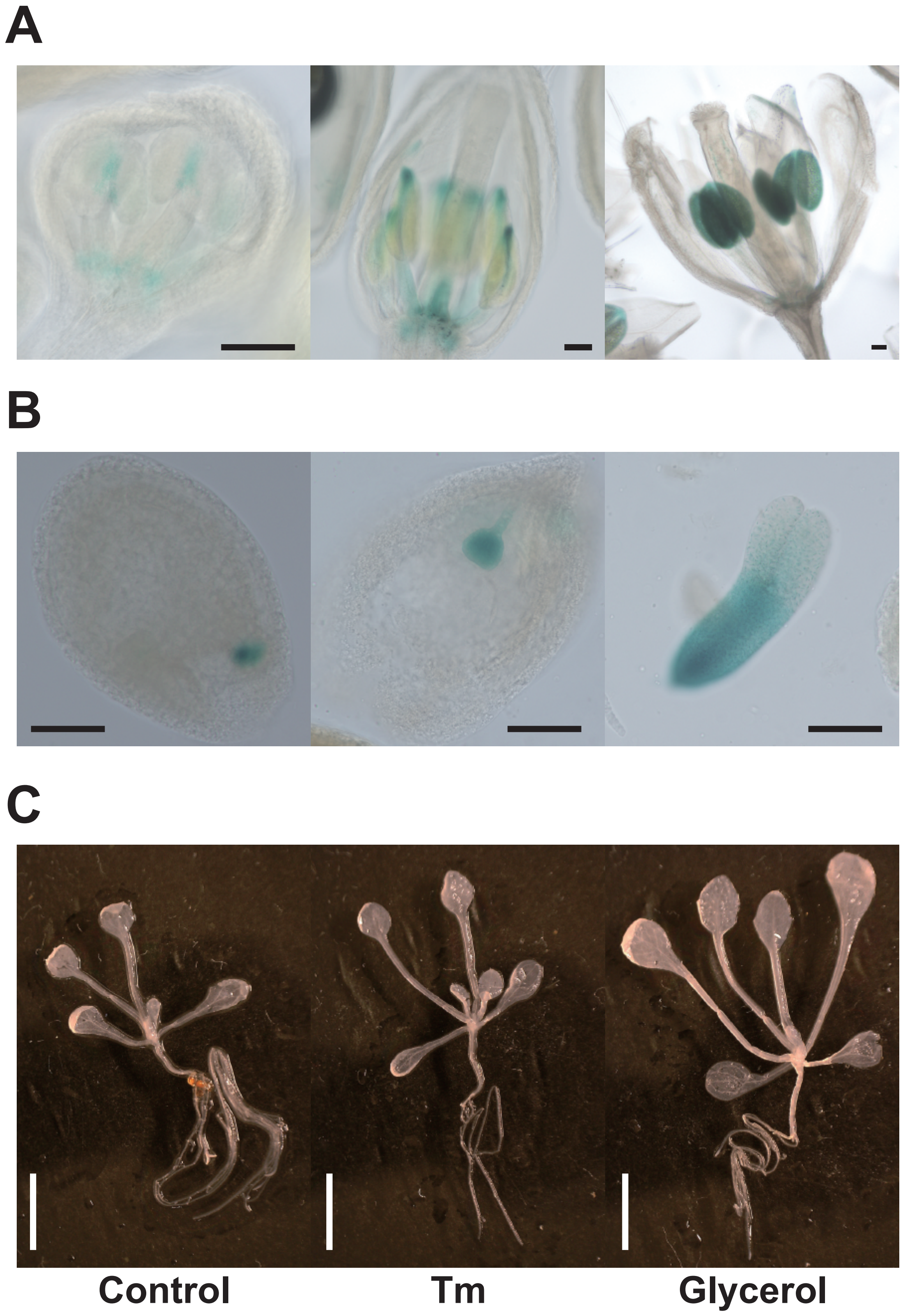


**Figure S3. Tissue-specific expression of *IRE1C* gene.** (A-C) GUS histochemical staining of transgenic Arabidopsis containing *IRE1C* promoter-*GUS* fusion construct in floral tissues (A), ovules and embryo (B). Bar = 100 μm. (C) 8-d-old seedlings treated with or without tunicamycin (Tm) for 5 h, and 11-d-old seedlings treated with glycerol for 3d. Bar = 5 mm.

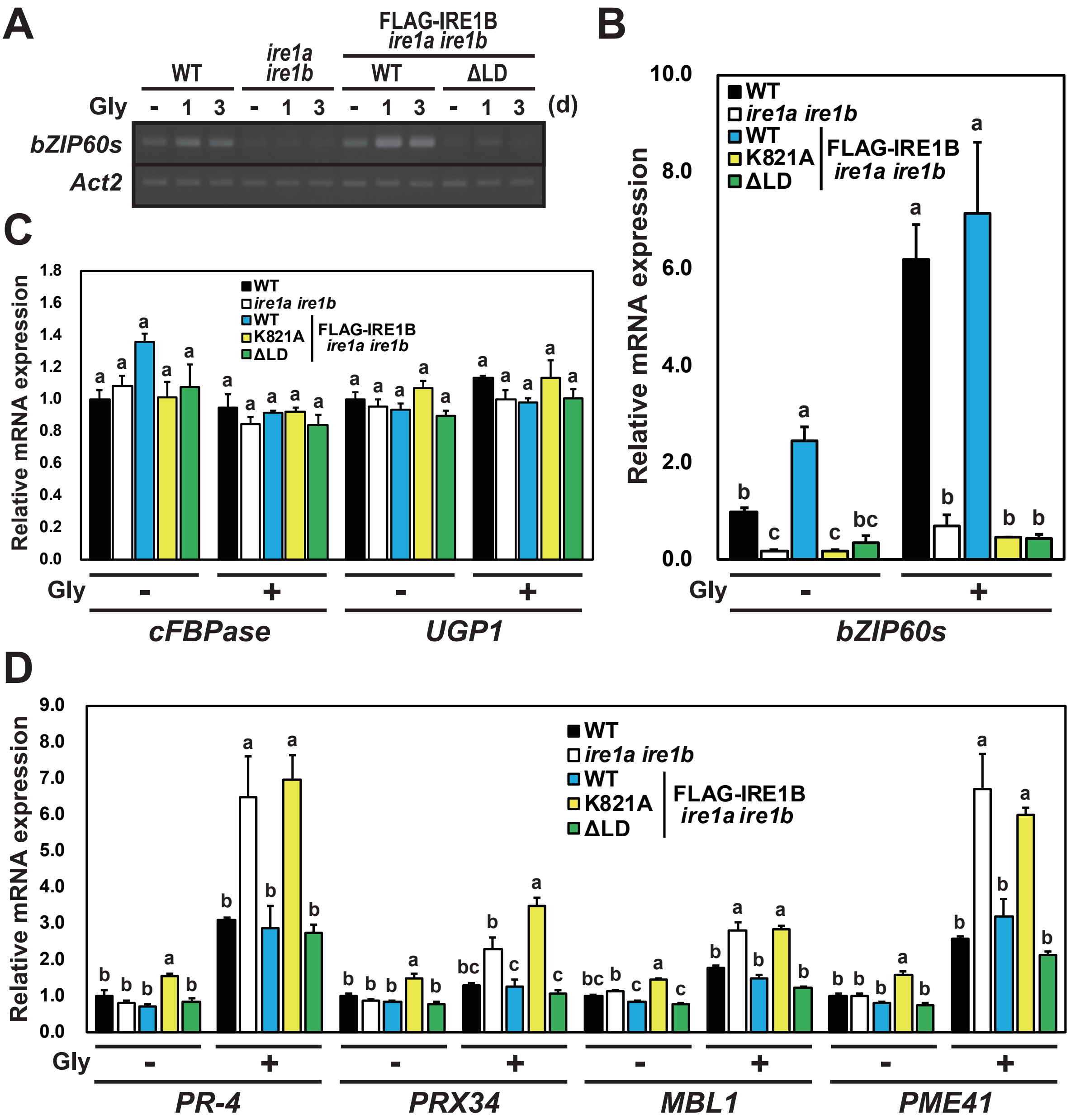


**Figure S4. Effect of RNase and sensor domain of IRE1B on *bZIP60* splicing and RIDD under glycerol treatment.** (A) Detection of *bZIP60s* mRNA splicing in WT, *ire1a ire1b*, and FLAG-IRE1B(WT, ΔLD) transgenic *ire1a ire1b* plants at 10 DAG. RT-PCR was performed using *bZIP60s*-specific primers. *Actin2* (*Act2*) was used as an internal control. Glycerol treatment was performed for 0, 1, and 3 d. (B-D) The relative mRNA levels of *bZIP60s* (B), cytosolic marker protein genes (C), and RIDD target genes (D) in WT, *ire1a ire1b*, and FLAG-IRE1B(WT, K821A, ΔLD) transgenic *ire1a ire1b* plants. RNA from seedlings at 10 DAG treated with (+) or without (-) glycerol for 3 d was subjected to qPCR. Data are means ± SEM of four independent experiments. Different letters within each treatment indicate significant differences ($P < 0.05$) by the Tukey–Kramer HSD test.

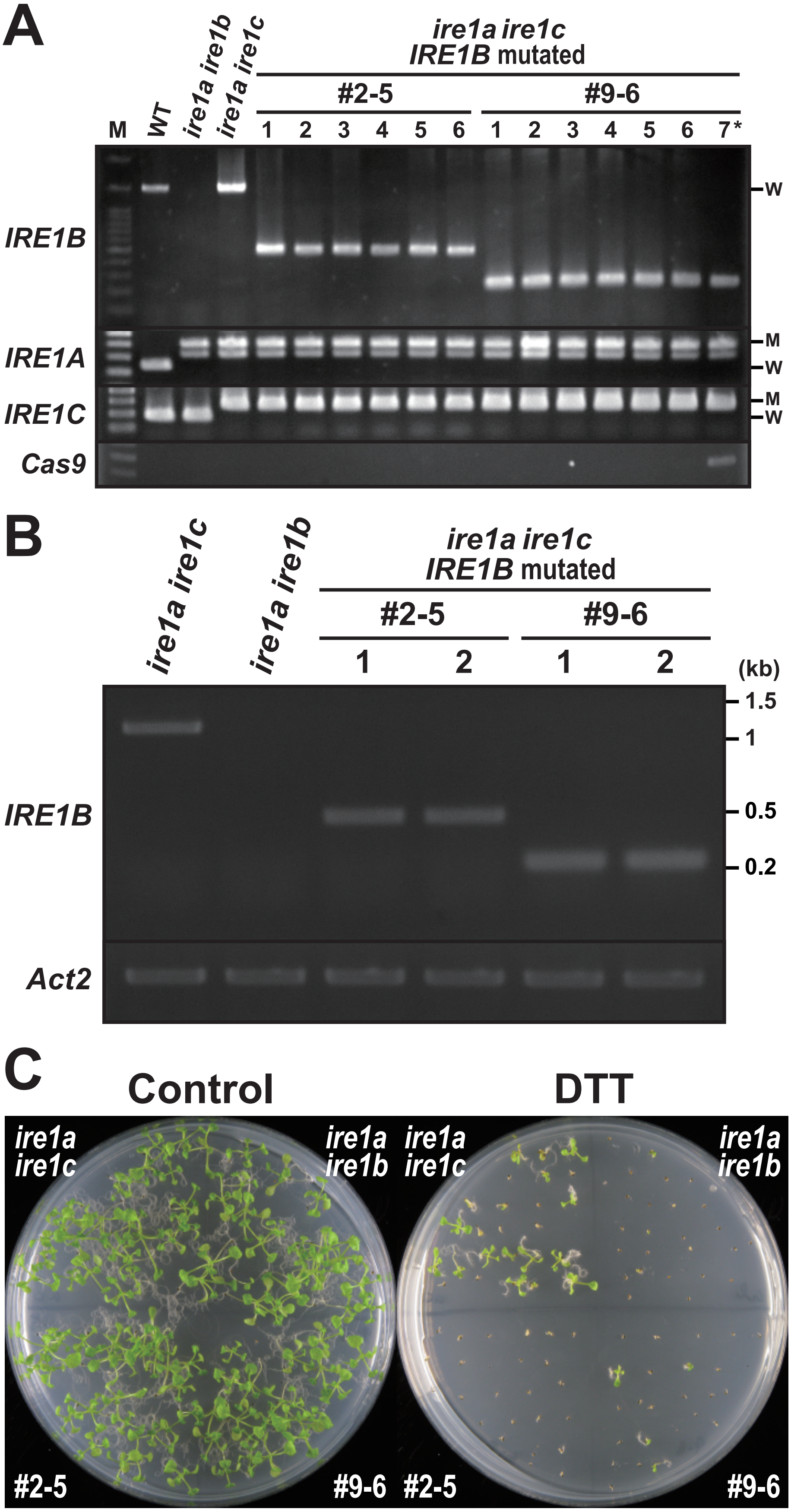


**Figure S5. Characteristics of the CRISPR/Cas9-mediated *IRE1B* mutant lines #2-5 and #9-6.** (A) Genotyping of *IRE1A-C* genes in wild-type, *ire1a ire1b*, *ire1a ire1c* mutants, and $T_3$ plants of lines #2-5 and #9-6. Lane M, 100 bp DNA ladder. Lanes 1-6, six independent plants for each line. M, mutant. W, wild-type. Note that PCR amplification of *Cas9* gene was performed to detect T-DNA, and that #9-6 lane 7 is a $T_2$ sibling plant having T-DNA used as a control. (B) RT-PCR of *IRE1B* mRNA in *ire1a ire1c*, *ire1a ire1b* mutants, and $T_3$ plants of #2-5 and #9-6. *Actin2* (*Act2*) was used as an internal control. (C) DTT sensitivity of the *ire1a ire1b*, *ire1a ire1c* mutants, and $T_3$ plants of #2-5 and #9-6. Seedlings at 15 DAG were treated with or without 1mM DTT.

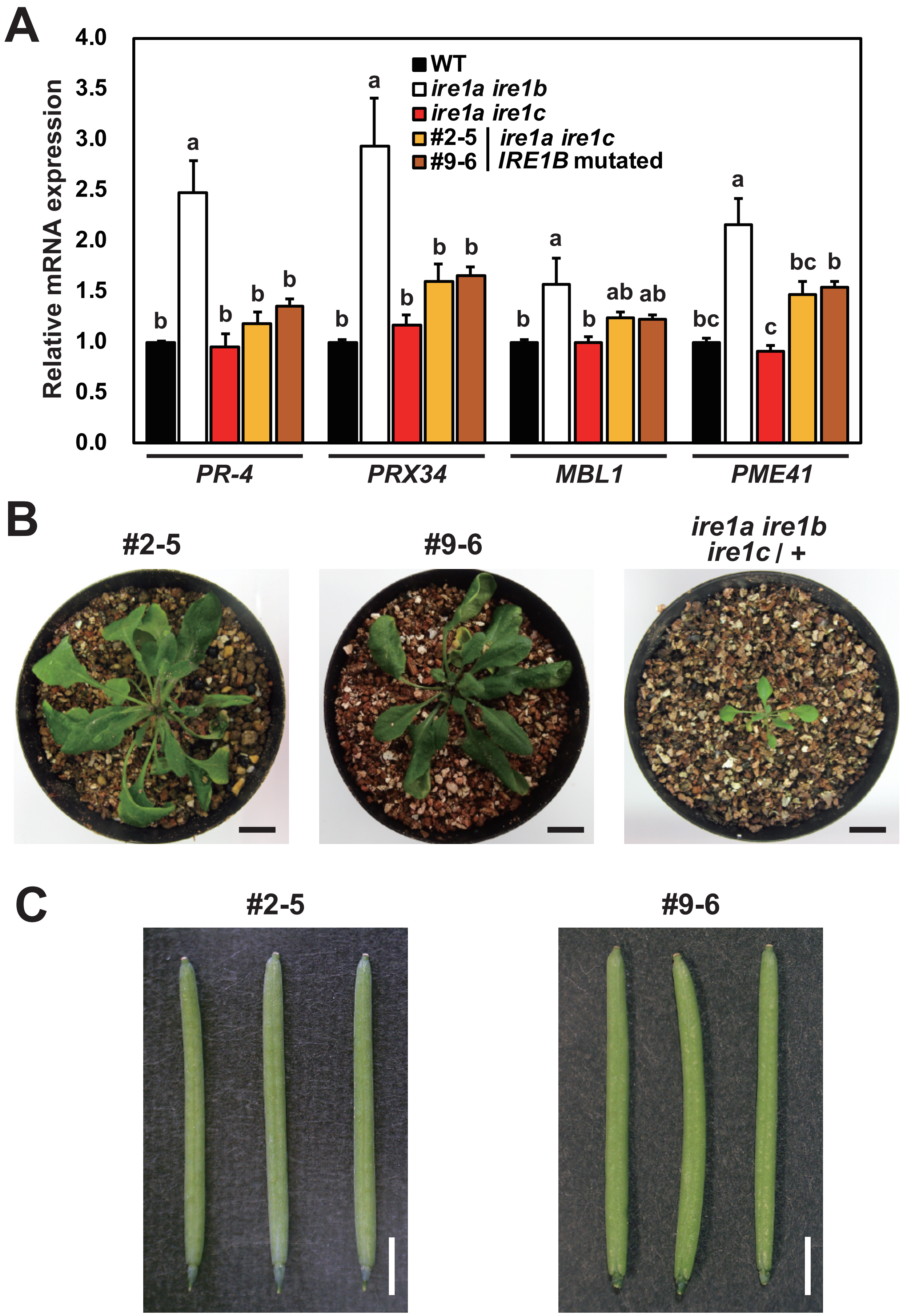


**Figure S6. #2-5 and #9-6 lines retain RIDD activity and show no phenotypic abnormality.** (A) The relative mRNA levels of RIDD target genes in WT, *ire1a ire1b*, *ire1a ire1c*, and *ire1a ire1c* with *IRE1B* mutation $T_3$ lines #2-5 and #9-6. RNA from seedlings at 10 DAG treated with glycerol for 3 d was subjected to qPCR. Data are means ± SEM of four independent experiments. Different letters indicate significant differences ($P < 0.05$) by the Tukey–Kramer HSD test. (B) $T_3$ plants of lines #2-5 and #9-6 and *ire1a ire1b ire1c*/+ mutant at 40 DAG. Bar = 10 mm. (C) Siliques of the lines #2-5 (left) and #9-6 (right) plants. Bar = 3 mm.
