## Supplemental Table 1 for "Unfolded protein-independent IRE1 activation contributes to multifaceted developmental processes in Arabidopsis"

**Table S1. Primers and oligonucleotides used in this study**

| Name | Sequence |
| --- | --- |
| Primers used for *ire1a* gene genotyping | |
| IRE1A(GT) FW  IRE1A(GT) RV  SALK(GT) RV | ATCGTCCACTTACACGAATTGGGAATAGTT  TTTTGCGAGATCAATCAGTCCTATTTTTGA  ACACTCAACCCTATCTCGGGCTATTCTTTT |
| Primers used for *ire1b* gene genotyping | |
| IRE1B(GT) FW  IRE1B(GT) RV  GABI(GT) FW | CAAAGTTTGGAGATGTTTTGTGGAGTGTTT  ATATCACAGTCTCAGGAAGAGGAAGGGAGA  TCTCCATATTGACCATCATACTCATTGCTG |
| Primers used for *ire1c* gene genotyping | |
| IRE1C(GT) FW  IRE1C(GT) RV  SALK(GT)2 RV | TGCTTACATTTGTCTAGAGCCTTGGAAATGCAG  TTAGCATTGGAGTTAGAGGTAATGGCTTCC  GCGTGGACCGCTTGCTGCAACTCTCTCAGG |
| Primers used for *BiP3* probe amplification | |
| BiP3 probe FW  BiP3 probe RV | ACAAACGAGATCGAAGAAGAGTTCTC  ACCGTCCCCAGTTTCTGCTCTTCGC |
| Primers used for *PR-4* probe amplification | |
| PR-4 probe FW  PR-4 probe RV | TCTGCTGCAGTCAGTACGGTTA  GCTGCATTGGTCCACTATTCTC |
| Primers used for *PRX34* probe amplification | |
| PRX34 probe FW  PRX34 probe RV | TATGCTCACCATTGCAGCTC  TGGTCGCTCTGGATAAGACC |
| Primers used for *MBL1* probe amplification | |
| MBL1 probe FW  MBL1 probe RV | GATTCTCCCACCGACACACT  CGTGTATGTCACGTCCCAAG |
| Primers used for RT-PCR of *bZIP60s* mRNA (Deng *et al*, 2011) | |
| bZIP60F4  b60SB2 | GAAGGAGACGATGATGCTGTGGCT  AGCAGGGAACCCAACAGCAGACT |
| Primers used for RT-PCR of *Act2* mRNA (Zhang *et al*, 2002) | |
| Act2 FW  Act2 RV | GTTGGTGATGAAGCACAATCCAAG  CTGGAACAAGACTTCTGGGCATCT |
| Primers used for qPCR of 18S rRNA (Zoschke *et al*, 2007) | |
| 25 5’ At18S  26 3’ At18S | AAACGGCTACCACATCCAAG  ACTCGAAAGAGCCCGGTATT |
| Primers used for qPCR of *bZIP60s* mRNA (Mishiba *et al*, 2013) | |
| bZIP60s-F-rt  bZIP60s-R-rt | AAGCAGGAGTCTGCTGTTGG  TTTGTGTGGGACATATAAGGGAAT |
| Primers used for qPCR of *PR-4* mRNA (Mishiba *et al*, 2013) | |
| PR-4-F-rt  PR-4-R-rt | GTGGGATGCTGATAAGCCGTA  TGCAGCATTTGTTCTTGTGTTCT |
| **Table S1.** (Continued) | |
| Name | Sequence |
| Primers used for qPCR of *PRX34* mRNA (Mishiba *et al*, 2013) | |
| PRX34-F-rt  PRX34-F-rt | ATGCGCAGATATGCTCACCA  AATGGAGCTGGAAGATTTGC |
| Primers used for qPCR of *MBL1* (AT1G78850) mRNA (Mishiba *et al*, 2013) | |
| AT1G78850-F-rt  AT1G78850-R-rt | CTTTGATTCTCCCACCGACA  CTTGGCTTCCATCACGAGAC |
| Primers used for qPCR of *PME41* (AT4G02330) mRNA | |
| PME41-F-rt  PME41-F-rt | GATGATTGGTGACGGAATAAACC  CGGTATTTCGGAAAGTCATGTTC |
| Primers used for qPCR of *cFBPase* (AT1G43670) mRNA | |
| cFBPase-F-rt  cFBPase-R-rt | ACGTTGGACCACACTGATGA  TTCCAGTGCTCAACACAAGC |
| Primers used for qPCR of *UGP1* (A3G03250) mRNA | |
| UGP1-F-rt  UGP1-R-rt | GTGAGAGTGAGAAGAGCGGATT  TTCTCGTAGGGAACAACGATTT |
| Primers used for *IRE1A* gene amplification | |
| IRE1A FW  IRE1A RV | CACCGTCTAGGACGCCTAGGCAC  CAGAAACGATGGATGTTTTCCCG |
| Primers used for *IRE1B* gene amplification | |
| IRE1B FW  IRE1B RV | CACCGTTGATACTCACGGAAGTCGG  GGGTACGGGTCTTTCAGATTG |
| Primers used for triple FLAG tag introduction into *IRE1A* gene | |
| IRE1A FLAG FW  IRE1A FLAG RV | GCCGACGATGTGACGTATCCGGACTACAAAGACCATGACGGTGATTATAAAGATCATGACATCGATTACAAGGATGACGATGACAAGATCGTTCCATCTAGTCCCGGCC  GGCCGGGACTAGATGGAACGATCTTGTCATCGTCATCCTTGTAATCGATGTCATGATCTTTATAATCACCGTCATGGTCTTTGTAGTCCGGATACGTCACATCGTCGGC |
| Primers used for triple FLAG tag introduction into *IRE1B* gene | |
| IRE1B FLAG FW  IREB FLAG RV | AAAGGATCTGAAATCTCCAAGGACTACAAAGACCATGACGGTGATTATAAAGATCATGACATCGATTACAAGGATGACGATGACAAGTTCTATGACAAATCTATCTCC  GGAGATAGATTTGTCATAGAACTTGTCATCGTCATCCTTGTAATCGATGTCATGATCTTTATAATCACCGTCATGGTCTTTGTAGTCCTTGGAGATTTCAGATCCTTT |
| Oligonucleotides used for gRNA constructs (underlines are the target sequences for *IRE1B* gene) | |
| gRNA1 linker FW  gRNA1 linker RV  gRNA2 linker FW  gRNA2 linker RV | GATTGATGGAAGCACGAAACAGGC  AAACGCCTGTTTCGTGCTTCCATC  GATTGTCGGATTGAGAGATTTGAT  AAACATCAAATCTCTCAATCCGAC |
| **Table S1.** (Continued) | |
| Name | Sequence |
| Primers used for *IRE1C* promoter amplification | |
| IRE1Cpro-FW  IRE1Cpro-RV | CACCGCAAAAAGCCAAAACATTTGA  AGCTTGGTCTCTTGAAGAATACAAAG |
